# Transmembrane channel-like 4 and 5 proteins at microvillar tips are potential ion channels and lipid scramblases

**DOI:** 10.1101/2024.08.22.609173

**Authors:** Seham Ebrahim, Angela Ballesteros, W. Sharon Zheng, Shounak Mukherjee, Gaizun Hu, Wei-Hsiang Weng, Jonathan S. Montgomery, Yaw Agyemang, Runjia Cui, Willy Sun, Evan Krystofiak, Mark P. Foster, Marcos Sotomayor, Bechara Kachar

## Abstract

Microvilli—membrane bound actin protrusions on the surface of epithelial cells—are sites of critical processes including absorption, secretion, and adhesion. Increasing evidence suggests microvilli are mechanosensitive, but underlying molecules and mechanisms remain unknown. Here, we localize transmembrane channel-like proteins 4 and 5 (TMC4 and 5) and calcium and integrin binding protein 3 (CIB3) to microvillar tips in intestinal epithelial cells, near glycocalyx insertion sites. We find that TMC5 colocalizes with CIB3 in cultured cells and that a TMC5 fragment forms a complex with CIB3 *in vitro*. Homology and AlphaFold2 models reveal a putative ion permeation pathway in TMC4 and 5, and molecular dynamics simulations predict both proteins can conduct ions and perform lipid scrambling. These findings raise the possibility that TMC4 and 5 interact with CIB3 at microvillar tips to form a mechanosensitive complex, akin to TMC1 and 2, and CIB2 and 3, within the mechanotransduction channel complex at the tips of inner ear stereocilia.

## Introduction

Microvilli are interconnected membrane bound actin-protrusions of uniform height that project from the apical surface of epithelial cells. Microvilli in intestinal epithelial cells (IECs) play critical functions including absorbing ions and nutrients, connecting with the glycocalyx and microbiome, and secreting vesicles containing antimicrobial peptides^1^. In addition to various chemical signals, microvilli are constantly subjected to mechanical stimuli due to peristalsis and sheer forces from luminal flow. Recent studies made the exciting discovery that microvilli possess mechanosensing ability^2,3^. However, the molecules and mechanisms that enable sensing, interpretation, and transmission of mechanochemical signals by microvilli are not well understood. Microvillar dysfunction is associated with a myriad of pathologies including microvillus inclusion disease^4–6^, celiac disease^7^, and Usher Syndrome type 1C^8,9^ underscoring the importance of addressing this knowledge gap.

Toward the distal tips of IEC microvilli resides a protocadherin-based intermicrovillar adhesion complex (IMAC) that connects neighboring microvilli. Within the IMAC, the extracellular domain of protocadherin-24 (PCDH24), also known as cadherin-related 2 (CDHR2), interacts with the extracellular domain of CDHR5 to form intermicrovillar links^10,11^, while their intracellular domains interact with Harmonin A (or USH1C)^11,12^, myosin 7^13,14^, and ANKS4^13,15^. Microvillar tips are also sites for secretion of glycolipids and, intriguingly, many protein components of the intestinal IMAC are shared with those forming the mechanotransduction apparatus in inner-ear sensory hair cells. Like IECs, hair cells feature apical actin-based membrane protrusions called stereocilia, which arrange in order of increasing height to form bundles that act as mechanosensory organelles. Similar to PCDH24 and CDHR5, cadherin-23 (CDH23) and protocadherin-15 (PCDH15) form oblique “tip links”-filaments connecting adjacent stereocilia in an arrangement that results in tensioning when bundles are sheared by mechanical stimuli from sound or head movements. This tension is conveyed to two members of the transmembrane channel-like (TMC) family of putative ion channels, TMC1 and 2, that localize to stereocilia distal tips^16,17^ and are the pore-forming subunits of the inner-ear mechanotransduction apparatus^18^.

The TMC proteins are evolutionary and structurally related to the mechanosensitive channels TMEM63/OSCA and the calcium-activated lipid scramblases or ion channels TMEM16 proteins^18–21^. TMC1 and 2 bind to calcium and integrin-bound (CIB) proteins 2 and 3^22–24^, and along with CDH23 and PCDH15, may form complexes with harmonin, myosin 7, and sans^13^ (or USH1G), a paralog of ANKS4. Many of these proteins are associated with Usher Syndrome, a disorder causing deaf-blindness as well as enteropathies^8,9^.

Here, using genetically modified mice and super-resolution fluorescence light microscopy, we find that two other members of the TMC family, TMC4, and TMC5, along with CIB3, localize to microvillar distal tips in IECs, near glycocalyx insertion sites. Moreover, co-localization of CIB3 and TMC4/5 in cultured cells and nuclear magnetic resonance (NMR) spectroscopy experiments suggest that CIB3 can directly bind to TMC4 and 5. Interestingly, TMC4 and 5 homology and AlphaFold 2 (AF2) models combined with molecular dynamics (MD) simulations suggest that these proteins are non-selective, low-conductance ion channels with possible lipid scramblase activity. These discoveries extend the molecular and structural similarities between microvilli and stereocilia and raise the exciting possibility that PCDH24, CDHR5, CIB3, TMC4, and TMC5 mediate mechanochemical signaling in IEC microvilli as CDH23, PCDH15, CIB2, CIB3, TMC1, and TMC2 do in inner-ear hair cell stereocilia. Whether intermicrovillar links formed by PCDH24 and CDHR5 or the glycocalyx directly activate the CIB3, TMC4, and TMC5 complexes remains to be determined.

## Results

### TMC4 and 5 localize to the intestinal epithelial brush border in humans and mice

The intestinal epithelium comprises a monolayer of IECs, including absorptive enterocytes and secretory cells (Fig. 1a). This monolayer is organized into two distinct levels: villi (Fig. 1a), which project into the lumen and are lined predominantly with mature absorptive enterocytes, and crypts (Fig. 1a), invaginations between adjacent villi where stem cells reside and constantly divide and differentiate. The apical surface of IECs is covered with microvilli (Fig. 1a) organized in precise arrays, known as the “brush border”, to increase the absorptive surface area. Adjacent microvilli are connected to each other by numerous lateral links (Fig. 1b). While TMC1 and 2 expression is most critical in the inner ear, TMC4 and 5 are more widely expressed in the body and enriched in enterocytes of the gastrointestinal tract^25,26^. To determine the localization of TMC4 and 5 proteins within human enterocytes, we acquired patient biopsy samples of healthy small intestine and performed immunofluorescence staining of cryo-sections using antibodies against TMC4 and 5. We found that TMC4 is strikingly enriched at the tips of enterocyte microvilli in human small intestine (Fig. 1c), while staining with the TMC5 antibody was unsuccessful. We then wanted to determine if this microvillar enrichment is conserved in mice. Neither antibody worked for immunofluorescence in mouse tissue, so we performed Western blots on purified murine brush border and found TMC4 and 5 to be enriched in this fraction (Fig. 1d). Based on these data, we decided to generate knock-in mouse models by tagging endogenous TMC4 with GFP, and endogenous TMC5 with mCherry (See Materials and Methods).

**Figure 1:**
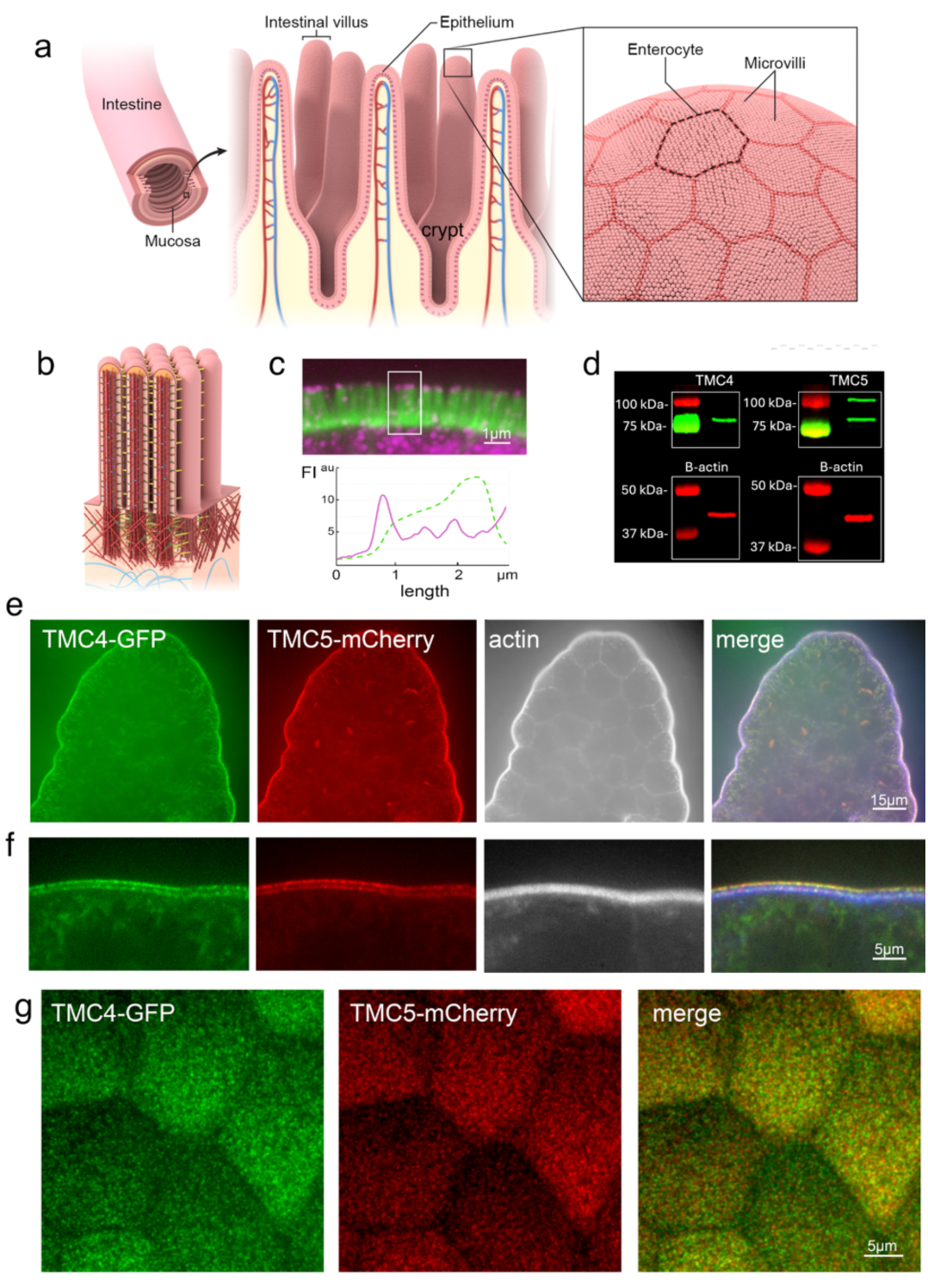
TMC4 and 5 localize to the distal tips and base of enterocyte microvilli. (a) Schematic showing the luminal side of the intestinal tract covered by a monolayer of epithelial cells, predominantly enterocytes, that are organized in repeating units of intestinal villi and crypts. The apical surface of each enterocyte is covered in arrays of microvilli-actin-based, membrane bound protrusions, known as the “brush border”. (b) Brush border microvilli are interconnected via several “lateral links”. (c) Confocal image of a cryosection of adult human small intestinal epithelium showing F-actin-based microvilli (green) and immunofluorescence staining of TMC4 (magenta), which is localized at the microvillar distal tips and also present in the cell cytoplasm. (d) Western blot of TMC4 and TMC5 on isolated mouse brush border. (e) A villus from the small intestine of a knockin mouse at P12 expressing TMC4-GFP (green) and TMC5-mCherry (red), stained with phalloidin to label F-actin (white). Both TMC4 and TMC5 are enriched at the apical surface of enterocytes. (f) TMC4-GFP (green) and TMC5-mCherry are localized to microvillar tips and base. (g) En face view of IECs from mouse expressing TMC4-GFP and TMC5-mCherry.

We visualized endogenous TMC4 and 5 in transgenic mice expressing TMC4-GFP and TMC5-mCherry and found that both are highly enriched at the brush border of murine IECs (Fig 1e), similar to TMC4 localization in human tissue (Fig. 1c). A cross-section view at high magnification reveals that both TMC4 and TMC5 localize specifically at the distal tips as well as the base of brush border microvilli (Fig 1f). From an en face view of the intestinal epithelial apical surface, we found that the TMC4 and 5 proteins are expressed robustly in microvilli across the cell surface (Fig. 1g). Additionally, we performed similar experiments in mCherry-TMC1 and GFP-TMC2 mice^16^ to examine whether TMC1 and 2 were also expressed in intestinal microvillar tips. As expected, these proteins were not detected in the intestinal microvilli (Fig S1).

### Microvillar localization of TMC4 and 5 shows spatio-temporal variation

From as early as postnatal day P1, robust GFP and mCherry fluorescence in the form of diffraction-limited puncta were observed at the distal tips of microvilli in all regions of the small intestine in mice expressing TMC4-GFP and TMC5-mCherry. The presence of discrete puncta, rather than a uniform diffuse fluorescence along the microvillar length, indicates an enrichment of a few closely associated TMC4 and 5 molecules in small clusters within each punctum (Fig. 2a). However, we often observed that both TMC4-GFP and TMC5-mCherry also localized along the microvilli as well as at a distinct region close to the microvillar base (Fig. 2a). We followed this expression pattern with age (Fig. 2a) and found that the relative localization of TMC4 and 5 at the microvillar base compared with the distal tip was reduced between P1 and P30. Conversely, in the proximal colon (Fig. 2b), TMC4 and 5 was enriched at the tips early postnatal (P2), and at both tip and base in adulthood. We also confirmed a strong expression of TMC4-GFP and TMC5-mCherry at the apical surface of IECs in intestinal crypts along the intestinal tract (Fig. 2c). We suggest that enrichment of TMC4 and 5 at the microvillar base may reflect the presence of a hub from which the TMC4 and 5 are transported to the distal tip.

**Figure 2:**
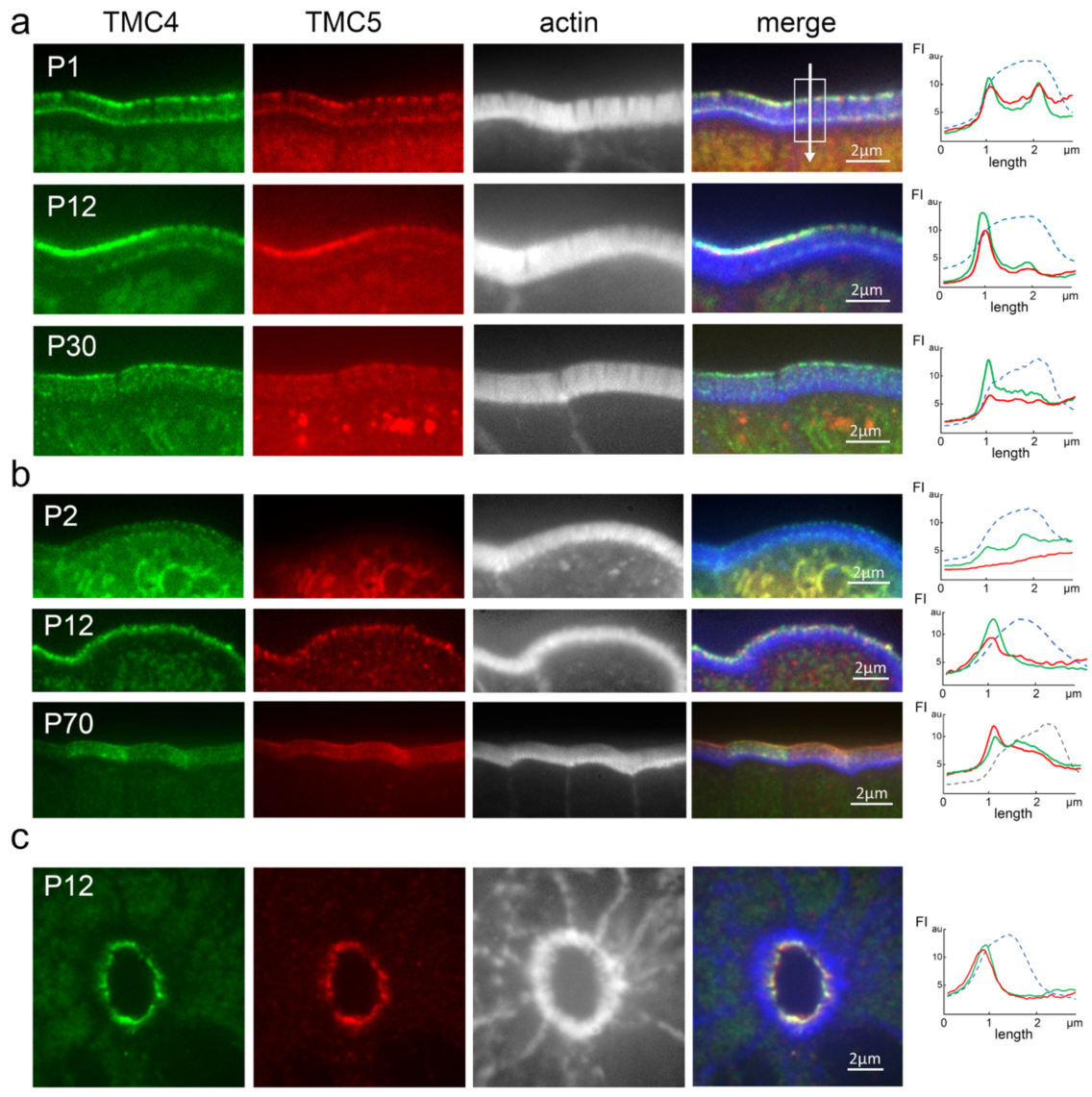
Microvillar localization of TMC4 and 5 shows spatio-temporal variation. (a and b) TMC4-GFP (green) and TMC5-mCherry (red) are enriched at the distal tips and base of microvilli (white) early postnatal (P1) in the small intestine (a), and only at microvillar tips in the large intestine (b). Enrichment along the microvilli varies with age in both intestinal segments. (c) TMC4-GFP (green) and TMC5-mCherry (red) are also enriched at the apical surface of enterocytes in intestinal crypts. Panels on right depict fluorescence intensity of TMC4-GFP (green), TMC5-mCherry (red) and actin (blue), along microvilli. The direction of the arrow in (a) depicts the direction of the *x*-axes, with the first fluorescence intensity (FI) peak corresponding to microvillar tip, and the second peak corresponding to the microvillar base.

### TMC4 and 5 localization at the distal tip of microvilli is spatially distinct from the microvillar Usher interactome

To assess the localization of TMC4 and 5 relative to known components of the microvillar IMAC, we labeled small intestinal tissue from mice expressing TMC4-GFP and TMC5-mCherry with antibodies against MYO7B, Harmonin A or ANKS4B (Fig. 3a). Fluorescence intensity line scans along the microvillar length showed that TMC4-GFP and TMC5-mCherry both localized consistently more tip-ward than all three Usher network proteins (Fig. 3a). These proteins have been previously referred to as microvilli “tip” proteins, however, our electron microscopy (EM) immunogold localization shows that they are excluded from the very tips of microvilli (Fig. 3c and d). This confirms the presence of two distinct compartments in the microvillar-tip region; one at the very distal tip, containing TMC4 and 5, and a second that is proximal to this, comprising ANKS4B, MYO7B and Harmonin A. The latter compartment is at the level of the cadherin-based lateral links which can clearly be seen by thin-section EM (Fig. 3b), and the imm unogold EM of Harmonin A (Fig. 3c) and MYO7B (Fig. 3d). We conclude from these analyses that most of the TMC4 an d 5 proteins are not directly associated with the Usher network proteins and with the lateral links.

**Figure 3:**
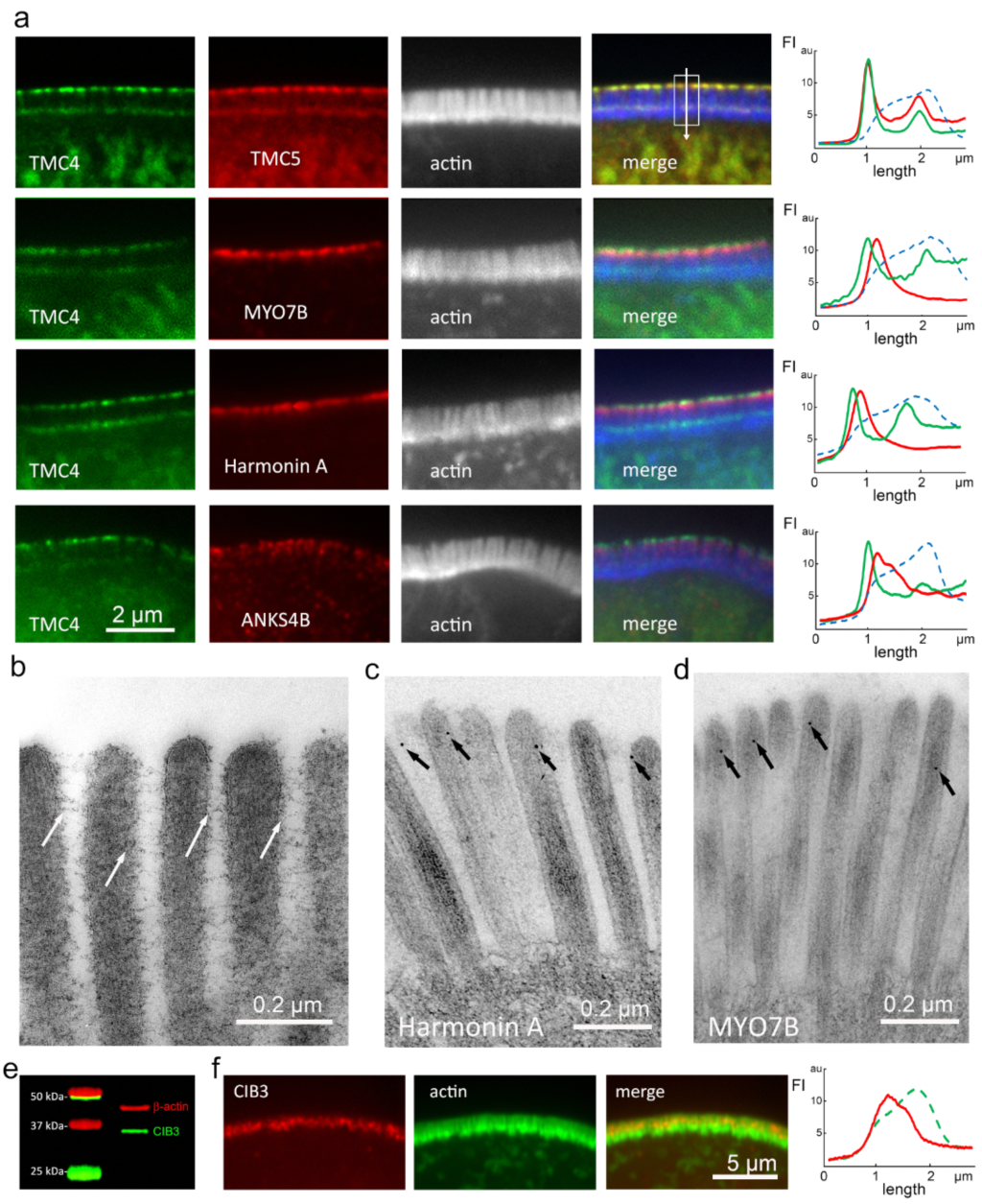
TMC4 and 5 localization at the distal tip of microvilli is spatially distinct from the microvillar Usher interactome. (a) Localization of TMC4-GFP (green) and TMC5-mCherry or various members of the IMAC (red) along microvilli (white) of small intestinal IECs from P12 knockin mice. Fluorescence intensity line scans along the microvillar length shown on the right (F-actin-blue, TMC4-green, TMC5 or IMAC proteins-red). The direction of the arrow in (a) depicts the direction of the *x*-axes, with the first FI peak corresponding to microvillar tip, and the second peak corresponding to the microvillar base. (b) Freeze-substitution thin section electron microscopy of microvilli from P4 mice showing inter-microvillar lateral links (white arrows). (c and d) Immuno-gold electron microscopy localization (black arrows) of Harmonin A (c) and MYO7B (d) in microvilli. (e) Western blot of CIB3 on mouse brush border. (f) Immunofluorescence staining of CIB3 (red) along microvilli (actin, green) of mouse enterocytes. Fluorescence intensity line scans along the microvillar length shown on the right.

### TMC4 and 5 expression levels exhibit cellular and microvillar-level variability

We next quantified the fluorescence intensity of TMC4-GFP and TMC5-mCherry in our knockin mice and observed a variability in the fluorescence intensity between adjacent cells (Fig. 4a), and also between neighboring microvilli in the same cell (Fig. 4b). Further, we found that while TMC4-GFP and TMC5-mCherry were often colocalized at microvilli tips they show variable stoichiometry, and sometimes showed distinct localizations (Fig. 4c and d). The distribution of fluorescence puncta seems to indicate that each microvillus tip contains either TMC4-GFP or TMC5-mCherry or both, suggesting that the two TMC proteins can be expressed as homo-or heterodimers. This is consistent with what was reported for TMC1 and 2 at the tips of stereocilia^16^. Freeze-etch electron microscopy of microvillar tips showed variable numbers of membrane proteins in this region (Fig. 4e), in line with the observation of variations in TMC4 and 5 expression levels in this region. Intriguingly, an inverse relationship between TMC4-GFP and TMC5-mCherry concentration at microvillar tips was also often observed (Fig. 4f), suggesting the two proteins may have both redundant as well as complementary functions.

**Figure 4:**
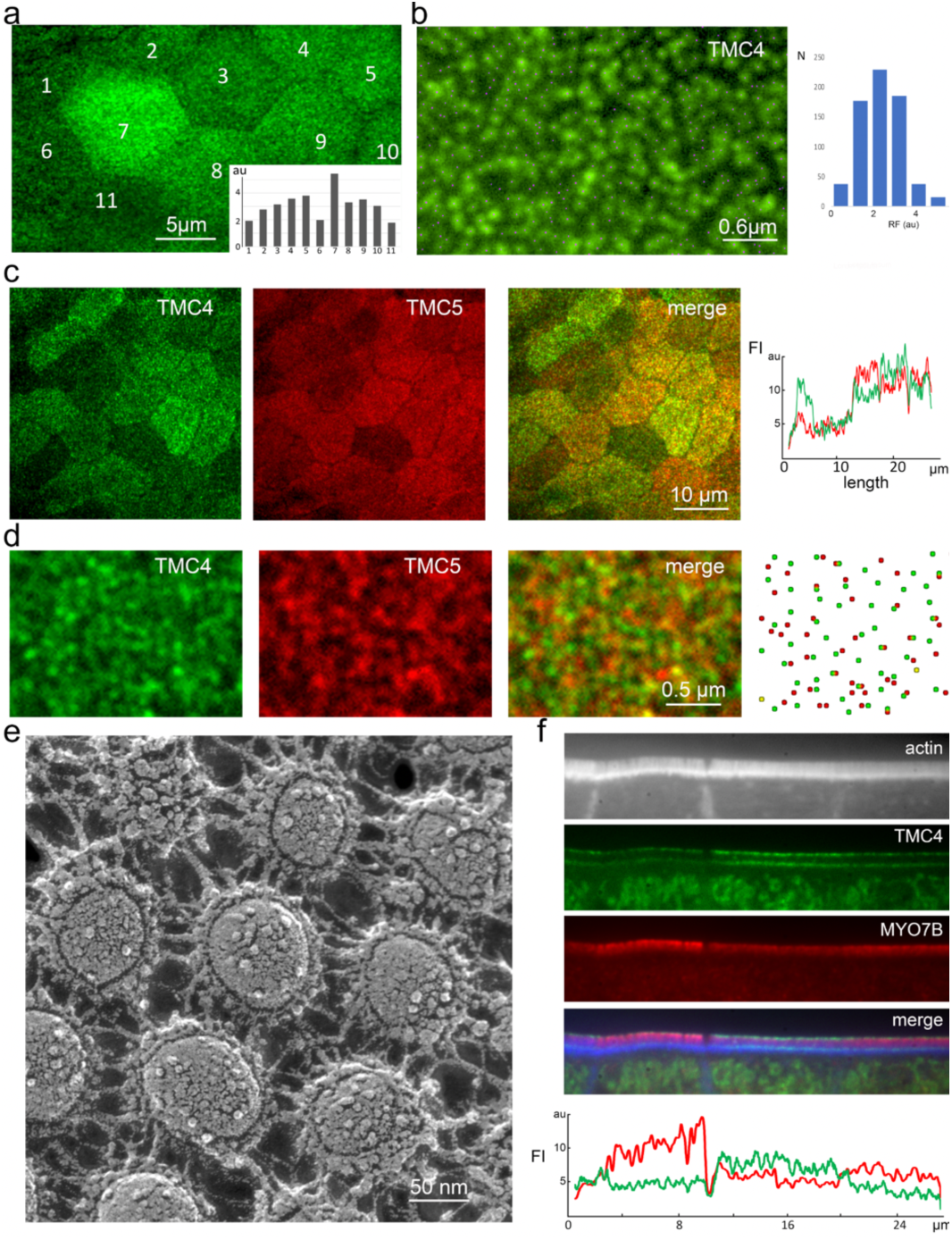
TMC4 and 5 expression levels exhibit cellular and microvillar-level variability. (a) En face view of enterocytes from knockin mouse at P12 expressing TMC4-GFP, with individual cells numbers, and average fluorescence intensity of GFP per cell plotted (inset). (b) TMC4-GFP at the tips of individual microvilli within a single cell localized by centroids (green dots), with fluorescence intensities plotted (right), RF= relative fluorescence and N = number of puncta. (c and d) Distribution of fluorescent TMC4-GFP (green) and TMC5-mCherry puncta show incidences of co-localization and also distinct localization. (e) Freeze-etch electron microscopy of microvillar tips from P4 mouse small IECs showing variable numbers of membrane proteins in this region. (f) Inverse relationship between TMC4-GFP (green) and TMC5-mCherry (red) concentration at microvillar tips (in FI) is often observed.

### TMC4 and 5 are enriched at the glycocalyx insertion site, which is subject to sheer forces

We next used freeze-etching electron microscopy to interrogate the local landscape of IEC microvillar tips in mouse tissue at the macromolecular level (Fig. 5). In this region, a rich glycocalyx network comprising highly diverse glycoproteins and glycolipids is secreted from the microvillar membrane. The glycocalyx is directly exposed to sheer forces from luminal contents and, in endothelial cells, has been identified to function as a mechanotransducer^27^. The components of the glycocalyx are covalently anchored and linked with the intracellular actin cortex via transmembrane proteins, thus enabling the transduction of extracellular signals to intracellular biochemical signaling pathways^27^. We observed inflections of the glycocalyx filaments at their connecting points at microvillar tips (Fig. 5e and inset), indicative of force-induced stretching or deformation. A surface view of the distribution of membrane-embedded proteins at a microvillar tip, where TMC4 and 5 are enriched, can be visualized when the fracture plane occurs precisely in this region, as seen in Fig. 5c and d. From our data, we thus posit that TMC4 and 5 localize to the microvillar membrane region near the glycocalyx (Fig. 5a and b) that experiences mechanical forces (Fig. 6).

**Figure 5:**
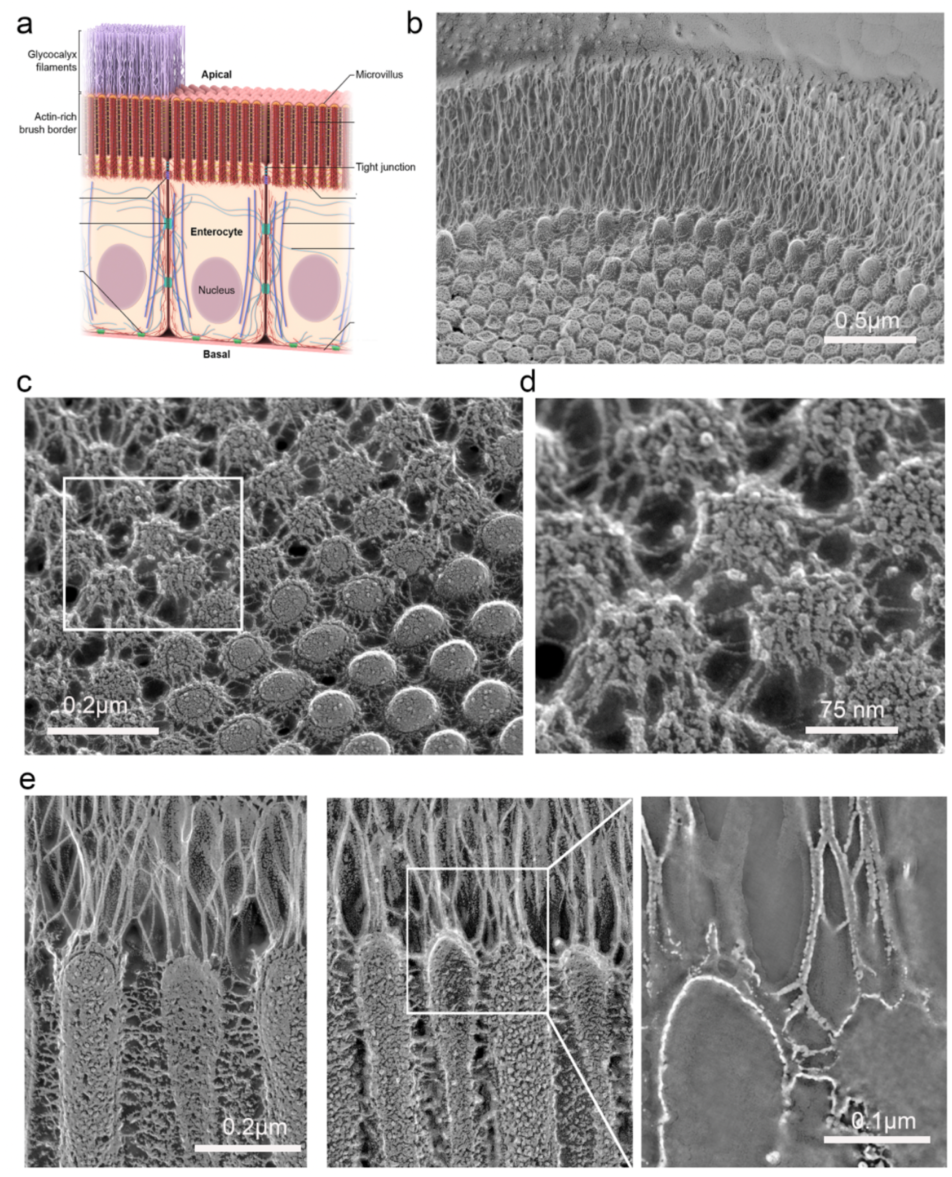
Microvillar tips are sites for glycocalyx insertion. (a) Schematic illustrating the glycocalyx network that is secreted from the tips of enterocyte microvilli. (b) Electron micrograph of a freeze-etch replica of mouse small intestine showing the stratified organization of the microvilli-rich brush border and glycocalyx layers. (c) Freeze-etching cross-fracture view of the microvilli tips at the point of insertion of the glycocalyx filaments on the membrane. (d) Close-up view of the area indicated by the rectangle in c showing the anchoring points of the glycocalyx filaments. (e) Left-The glycocalyx filaments emerging from the tips of the microvilli are distinct from the lateral links between microvilli. Right and inset-inflections of the glycocalyx filaments are observed at their connecting points to the microvillar membrane.

**Figure 6:**
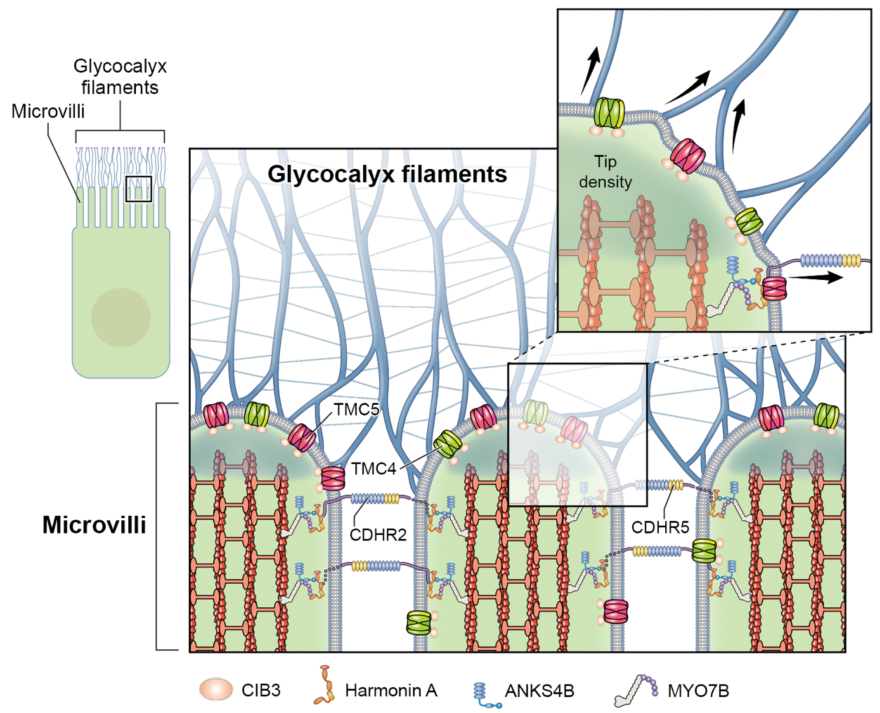
Diagram illustrating the localization of the TMC4-TMC5-CIB3 complex at the tips if IEC microvilli.

### CIB3 is also enriched at microvillar tips and colocalizes with TMC5 in a heterologous cell line

CIB2 and CIB3 were recently found to be integral subunits of the mechanotransduction channel in stereocilia^23^. Genetic, biochemical and structural studies have confirmed that CIB2/3 interact with TMC1 and 2^22–24,28^. The first intracellular loop (IL1) region of TMC1 and TMC2 interacts with CIBs and has a sequence motif conserved across TMCs and involved in the interaction (Fig. S2). Therefore, we investigated whether CIB2 and 3 are also present in intestinal microvilli and could potentially be part of a protein complex containing TMC4 or TMC5. Using Western blot in a mouse brush border fraction, we found that CIB3 is in fact present in intestinal epithelial cell microvilli (Fig. 3e). We then validated the enrichment of CIB3 in the microvilli by immunofluorescence staining and discovered that it localizes toward the distal tips (Fig. 3f). Finally, we tested whether CIB3 interacts with TMC4 and/or TMC5 by co-transfecting cDNA encoding each protein in COS7 cells, and determining the extent of co-localization, as we have done previously^13^. As seen with other TMC family proteins, we found that TMC4 and TMC5 are ensnared within the endoplasmic reticulum (ER) in COS7 cells (Fig. S3). CIB3 remains diffuse in the cytoplasm when overexpressed in COS7 cells (Fig. S3a), but when co-transfected with either TMC4 or TMC5, CIB3 localizes to the ER (Fig. S3) as it is binding to the TMC4 or TMC5. Together these data support the direct interaction of CIB3 with TMC4 and TMC5 at IEC microvillar tips, synonymous with CIB2 and CIB3 interactions with TMC1 and TMC2 at stereocilia tips.

### *In-vitro* interactions of TMC5 fragments with CIB3 suggest complex formation

To determine whether CIB3 directly interacts with TMC5 *in vitro*, we performed NMR experiments that monitor changes in amide resonances upon reconstitution of *Homo sapiens* (*hs*) CIB3 with *Mus musculus* (*mm*) TMC5-IL1 (equivalent to *hs* TMC5-IL1 T566A). Uniformly ^15^N labeled [U-^15^N] *hs* CIB3 was co-refolded in the presence of unlabeled *hs* TMC5-IL1 T566A and the complex was purified by size exclusion chromatography (SEC) in which a single peak was observed (Fig. S4a). TROSY ^1^H-^15^N correlation spectra were collected for the co-refolded mixture of [U-^15^N] *hs* CIB3 + *hs* TMC5-IL1 for comparison with the spectrum of [U-^15^N] *hs* CIB3 alone^22^ (Fig. S4b). We observed extensive chemical shift perturbations and the appearance of new peaks, and fewer broad peaks, consistent with binding-coupled folding^29–31^. A spectrum of [U-^15^N] *hs* CIB3 prepared with limited *hs* TMC5-IL1 (3:1 CIB3:TMC5-IL1 ratio) had peaks that overlay with both the apo and bound peaks, indicating a tight and stable interaction between CIB3 and TMC5-IL1 (Fig. S4c). These results suggest that family-wide interactions between TMCs and CIB2 and 3 are possible.

### Homology-based TMC4 and 5 structural models

The TMC proteins are evolutionary and structurally related to the TMEM16 and TMEM63/OSCA families of proteins. Cryo-EM and crystal structures of several members of the TMEM16 and TMEM63/OSCA proteins have revealed that these related proteins share a similar overall fold despite their low sequence identity^18–20,32^. These data suggest that TMC proteins would also present a similar structure, as now confirmed by structures of the invertebrate TMC-1 and TMC-2^28,33^. To gain insight into the pore properties of TMC4 and TMC5, we built homology-based structural models for monomeric murine TMC4 and TMC5 (Fig. 7a-c). Out of all the structures available of TMEM16 and OSCA proteins, a model based on the structure of the *Aspergillus fumigatus* (*af*) TMEM16 (PDB ID: 6E10) was the best template for TMC5 and the structure of the *Nectria hematococca* (*nh*) TMEM16 protein (PDB ID:4WIT) was the best template to model the structure of TMC4 (see Material and Methods section). These structural models revealed the presence of 10 transmembrane (TM) helices and a large pore at the protein-lipid interface formed by TM4-TM8 like that observed in the template structures, prior TMC1 models^19,34,35^, and recent TMC-1 and TMC-2 structures^28,33^.

**Figure 7:**
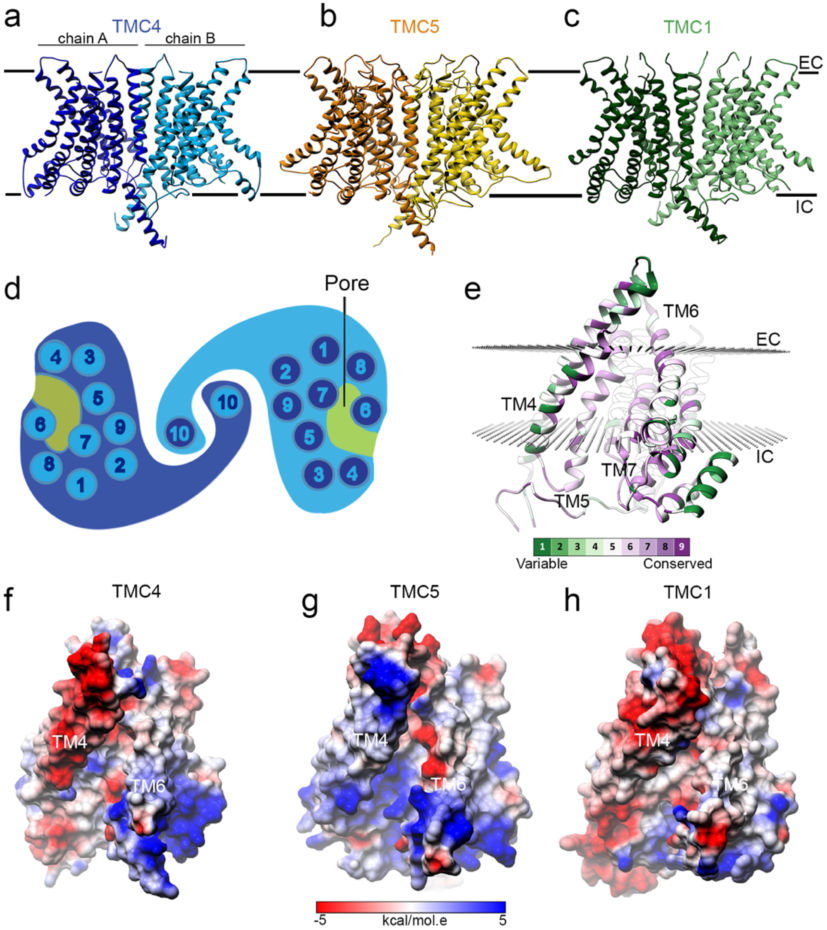
Structural homology models for TMC4 and 5. **(a-c) Ri**bbon representation for the TMEM16-based models of TMC4 (a), TMC5 (b), and TMC1 (c). Chains A and B are indicated and colored in different tones. Two black lines indicate the approximated localization of the plasma membrane with intracellular (IC) and extracellular (EC) locations indicated. (d) TMC topology. (e) Schematic representation of TMC transmembrane helices and pore. (f) Conservation of the TMC residues lining the pore estimated by ConSurf. TM4-7 are indicated. Two spheres of planes indicate the approximated localization of the membrane bilayer calculated by OPM. (g-i) The charge surface at the pore of TMC4 (g), TMC5 (h), and TMC1 (i) depicts the electrostatic differences in this region. TM4 and TM6 lining the pore are indicated.

The pore cavity is highly conserved between TMC proteins suggesting an important role for this structural feature in the function of these proteins. The TM4-TM8 pore represents an open conformation like that observed in the structures used as templates (Fig. 7d-e). We looked at the conservation of the properties of the residues that built this cavity (Fig. 7f) and found that the cavities of TMC4 and TMC5 present less negatively and more positively charged residues than TMC1 leading to a more positive cavity (Fig. 7g-i).

### AF2-based TMC4 and 5 structural models

Recent advances in protein structure prediction also offer the possibility to create accurate models of proteins complexes without templates^36^. To complement our homology modeling, we used AF2 to create additional models of TMC4 and 5. Since there is experimental evidence that TMC1 assembles as a dimer^28^ and structures for *nh* TMEM16^35^, *mm* TMEM16A^37^, *af* TMEM16^34^, TMEM16K^38^, TMEM16F^39^, *ce* TMC-1^28^, and *ce* TMC-2^33^ are all dimeric, we used AF2^36^ and AF2 multimer^40^ to generate models of mouse TMC4 and 5 dimers. The dimeric interface is formed by swapped TM10 helices as observed in the TMC-1 and TMC-2 structures (Fig. 8a-c). As expected, the putative pore is lined up by helices TM4 through TM8, with TMC4 and 5 cavities featuring less negatively charged residues when compared to TMC1 and TMC2. The monomeric subunits are similar to our homology-based models with ten transmembrane helices (Fig. 8). Unlike our homology-based models, the ion conduction pathway appears occluded by several residues stemming from TM4 and TM6 and interacting with each other (Fig. S5a,c).

**Figure 8:**
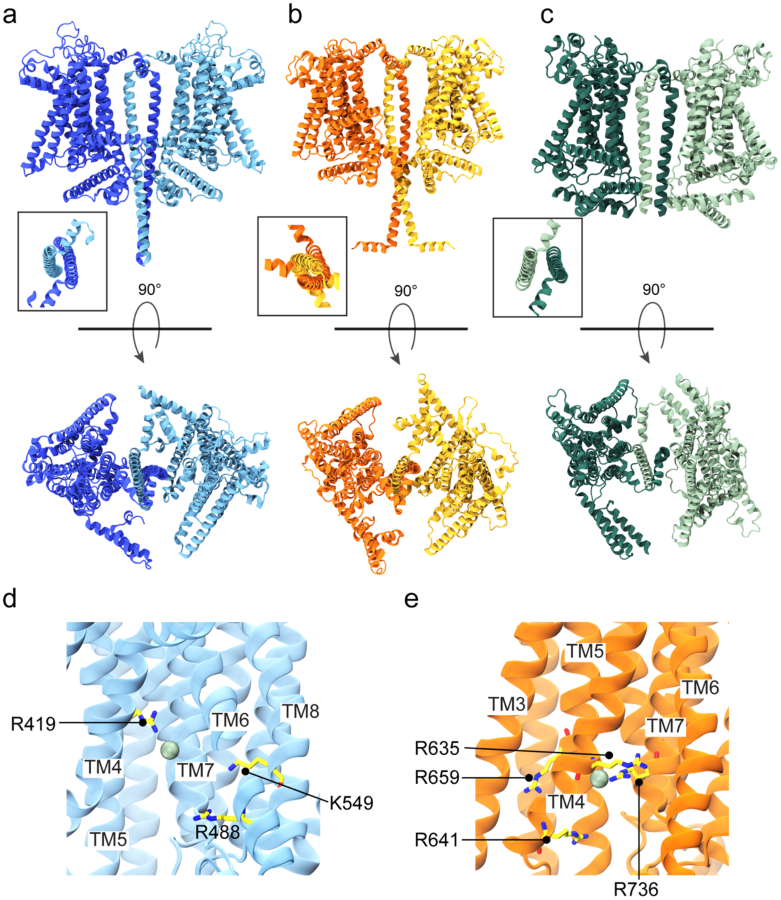
AF2-based structural models for TMC4 and 5. (a-b) AF2-based *mm* TMC4 and *mm* TMC5 dimers with side (top) and top-down (bottom) views. (c) Side and top views of dimeric *ce* TMC-1 cryo-EM structure. Insets show TM10 swaps in each system. (d-e) Cl^-^ binding residues in TMC4 and TMC5, respectively.

### MD simulations of TMC4 and 5 reveal a hydrated closed pore and a dynamic Cl^-^ binding site

To test the ion-conduction properties of TMC4 and 5 we carried out all-atom MD simulations of our AF2 models embedded in a hydrated POPC bilayer with 300 mM NaCl (Table S1). Equilibrium 100-ns long simulations revealed quick hydration of the putative pore from both extracellular and intracellular sides, but there was a discontinuity in the middle as seen in water density maps (Fig. S6a,c). Average electrostatic maps (Fig. S6e,g) indicated that the cavity for these proteins were more positively charged compared to the bulk, consistent with distribution of charges and a possible non-selective or anion selectivity. We also performed MD simulations with voltage^41^ starting from the end of the equilibration runs. Voltage was set to ± 0.5 V for TMC4 and 5. During the equilibrations and simulations with an applied negative voltage we observed Cl^-^ ions entering the channels from the cytoplasmic side, but crossings were not observed under any simulation conditions. In TMC4, the Cl^-^ ion at the pore interacted with and oscillated between three positively charged residues: R419, R488, and R635 (Fig. 8d; Movie 1). In TMC5, the Cl^-^ ion interacted mostly with R641 and R736, but also with R659 and K549 (Fig. 8e). Overall, these simulations show possible paths of ion permeation for TMC4 and 5, but the absence of crossing events suggest that our AF2 models do in fact feature closed pores.

### MD simulations of TMC4 and 5 with open pore from homology modeling show ion conduction

Although our AF2 models of TMC4 and 5 likely represent closed channels, the homology-based models feature wider pores (Fig. S5b,e). Hence, the pore regions from our TMC4 and 5 homology-based models were aligned to one of the monomers in the corresponding AF2 models and used as targets for open pores in targeted MD (TMD) simulations^42^. This resulted in TM4 and TM6 moving away from each other and the formation of wider pores that quickly got hydrated (Movie 2). The other non-targeted monomers within the TMC4 and 5 dimers were kept unaltered as controls. After TMD, we locked the open conformations of the targeted monomers by either fixing or constraining backbone atoms of the pore-forming residues (320 to 560 in TMC4 and 560 to 780 in TMC5) in 100-ns long equilibrations followed up by 200 ns of dynamics with applied voltage (± 0.5 V). The 100-ns long equilibrations showed intact water channels for the targeted monomers (Fig. S6b,d). Interestingly, POPC headgroups inserted themselves into the hydrophilic cavity harboring the water channels and we observed multiple Na^+^ and Cl^-^ ions crossing the TMC4 and 5 channels in opposite directions in simulations at -0.5 V (Table S2; Movie 3). In general, the number of Cl^-^ ions at the cytoplasmic mouth of the pores was larger than the number of Cl^-^ ion crossings under voltage, as these ions interacted with the same positive residues highlighted in equilibrium simulations, namely R419, R488, and R635 in TMC4 and R641, K549, R659, and R736 in TMC5.

Ion crossings in simulations of TMC4 post-TMD at +0.5 V were minimal. In contrast, the 200-ns long simulation of TMC4 post-TMD at -0.5 V with a fixed open conformation for one monomer showed a total of 17 Na^+^ and 3 Cl^-^ ion crossings with an estimated upper limit for the conductance of α = 32 pS (Fig. 9a and Table S2). Conductance values for two other independent simulations at -0.5 V with the open conformation constrained for one monomer were α = 9.6 pS (6 Na^+^ and 0 Cl^-^) and α = 25.6 pS (14 Na^+^ and 2 Cl^-^). The estimated conductance in a third simulation performed using the Anton2 supercomputer and lasting 480 ns at -0.5 V was α = 24.7 pS (29 Na^+^ and 8 Cl^-^; Fig. 9 and S7). In simulations at -0.5 V where multiple crossing events were observed we noticed that Na^+^ ion crossings correlated with the presence of one or more Cl^-^ ions in the cytoplasmic mouth of the pore (Fig. S8a-c), suggesting that TMC4 might be an anion-assisted non-selective ion channel.

**Figure 9.**
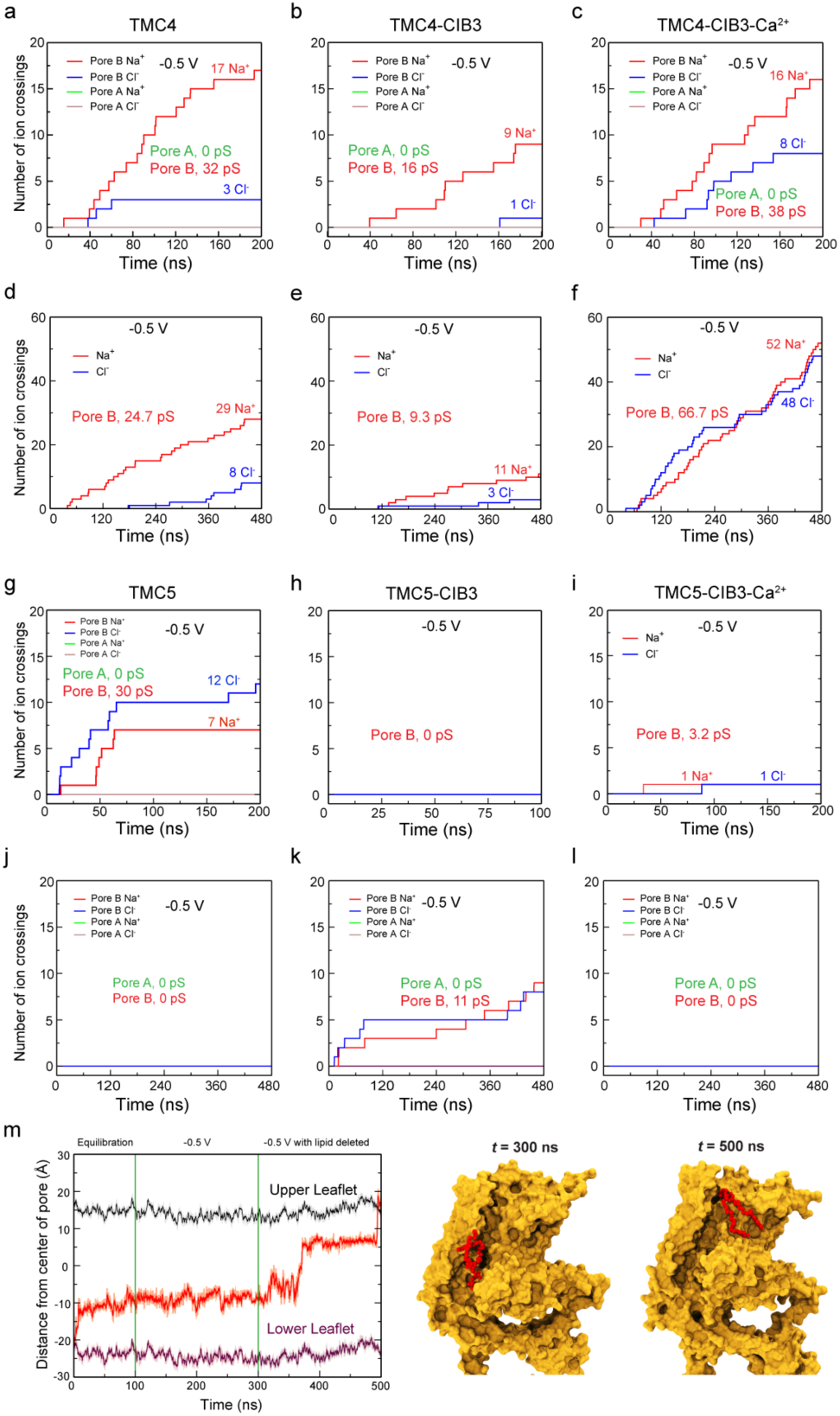
Ion conduction and lipid scrambling in TMC4 and 5 proteins. (a-f) Number of ion crossings vs time for TMC4 (a,d), TMC4-CIB3 (b,e), and TMC4-CIB3-Ca^2+^ (c,f) carried out at short (200 ns; a-c) and longer (480 ns – Anton2; d-f) timescales using -0.5 V. (g-l) Number of ion crossing vs time for TMC5 (g,j), TMC5-CIB3 (h,k), and TMC5-CIB3-Ca^2+^ (i,l) carried out at short (200 ns; g-i) and longer (480 ns – Anton2; j-l) timescales using -0.5 V. In all these cases monomer A is in closed conformation and monomer B was opened using TMD. (m) Full trajectory of a flipping lipid (555) from equilibration to the end of a voltage simulation of TMC5 system. Snapshots of lipid conformations are shown at indicated times. Some panels are also in Fig. S7.

Analyses of simulations for TMC5 post-TMD revealed Na^+^ and Cl^-^ ions visiting the pore but no conduction of ions. This was caused by a lipid blocking the channel. We removed the lipid and continued simulations at - 0.5 V, which revealed conduction with a slight preference of Cl^-^ over Na^+^ ions (Fig. 9g). Multiple lipid molecules from the cytoplasmic side lined up the channel and 60 ns after removal of the blocking lipid, another lipid inserted itself deep inside the channel and switched to the extracellular leaflet of the membrane (Fig. 9m and S8d; Movie 4). Although ions were in the vicinity of pore-lining residues, this lipid scrambling event led to a blockage of ion conduction. The self-insertion of lipids into the hydrophilic cavity of TMC4 and the lipid scrambling event in TMC5 suggest that apart from being putative ion channels, TMC4 and TMC5 might function as scramblases, a prediction that remains to be tested experimentally.

### Sequence analyses and simulations highlight residues involved in ion selectivity

A multiple sequence alignment (Data S1) for human and mouse TMC1, 2, 4, and 5 allowed us to identify residues potentially important for selectivity. In our TMC4 simulations, entrance of a Na^+^ ion from the extracellular side was facilitated by residues E454, E353, and E213. The ion then interacted with residues D375 (TM4) and E465 (TM6). Residue E353 is conserved in TMC5 but not in TMC1 and 2, residue E454 is not conserved in TMC1, 2, and 5, and residue E213 is conserved in TMC2, but not in TMC1 or 5. However, there is a nearby aspartate residue in TMC1 that might function similarly. Residue D375 is not conserved in TMC1, 2, and 5 but residue E465 is conserved. Towards the intracellular side, Na^+^ ions were coordinated by residues D473 (TM6) and D512 (TM7). Residue D473 is conserved across all four TMCs while residue D512 is exclusive to TMC4. Residues E465 and D473 are equivalent to E520 and D528 in mouse TMC1. Mutations of these residues in TMC1 results in decreased conductance^43^, consistent with our simulation predictions.

We also identified residues interacting with Cl^-^ ions during our simulations. Residue R419 and K549 in TMC4 are conserved in TMC5 (K664 and K773 respectively) but not in TMC1 and 2, whereas residue R488 (R717 in TMC5) is conserved across TMC1, 2, 4, and 5. Residues R419 and K549 might be responsible for Cl^-^ permeation in the channel. TMC5 has more positively charged residues when compared to TMC4: In addition to K664, K773 and R717, TMC5 has residues R635, R641, R736 facing the hydrophilic cavity, none of which are conserved in TMC1, 2, and 4. Interestingly, R736 in TMC5 is equivalent to D512 in TMC4 which might explain the higher Cl^-^ permeation or selectivity in TMC5 compared to TMC4 in our simulations.

### *In-silico* interaction of TMC4 and 5 with CIB3

Given our data showing potential interactions between TMC4 and 5 and CIB3 (Figs S3 and S4), we generated AF2 models for TMC4 and 5 in complex with CIB3. As observed in the structure of mouse TMC1-IL1 interacting with human CIB3 (Fig. S2; PDB:6WUD)^23^, our AF2 models show the mouse TMC4- and 5-IL1 fragments similarly interacting with mouse CIB3 (Fig. S9). Several residues at the binding interface are conserved across members of the TMC family, supporting the notion that CIB3 might be interacting with TMC4 and TMC5 in microvillar tips.

We used our AF2 models of the complexes to carry out additional MD simulations testing how CIB3 may alter TMC function. CIB proteins have two stable Ca^2+^-binding sites. Thus, two different systems were prepared, one each for TMC4 and 5 with CIB3 and TMC4 and 5 with CIB3 and bound Ca^2+^. These systems were placed in a hydrated POPC bilayer with 300 mM NaCl. No ion crossings were observed in equilibrations and simulations with applied voltage. As before, TMD simulations were used to open the pores followed by backbone constrained equilibrations and simulations with applied voltage.

Analyses of simulations of the post-TMD TMC4-CIB3 system show few ion crossings, despite its wider pore (Fig. S5c). Interestingly, in three out of four simulations of post-TMD TMC4-CIB3-Ca^2+^ systems at -0.5 V, we monitored more Cl^-^ ion crossings compared to what was seen in the post-TMD TMC4 and TMC4-CIB3 systems (Fig. S7), suggesting that Ca^2+^ bound to CIB3 enhances Cl^-^ conduction (Movie 5). Simulations using Anton2 and lasting 480 ns revealed more crossings and an estimated conductance of α = 67 pS. In contrast, we observed very few or no ion crossings in simulations at +0.5 V for the TMC4-CIB3 and TMC4-CIB3-Ca^2+^ systems. Overall, the pore of post-TMD TMC4-CIB3-Ca^2+^ was not selective for anions over cations or vice versa.

Analyses of simulations of the post-TMD TMC5-CIB3 and TMC5-CIB3-Ca^2+^ systems show very few ion crossing events, likely because of lipids inserting into the TMC5 pore cavity. In one of the simulations for TMC5-CIB3 at -0.5 V we see a lipid starting to flip from one side of the membrane to the other, similar to what we observed for TMC5 alone. Although further experiments and longer atomistic simulations are required to explicitly study the role of CIB3 in TMC4 and 5 function, our simulations suggest that upon binding and depending on whether Ca^2+^ is bound to it, CIB3 may alter the pore properties of TMC4 and 5.

## Discussion

Microvilli enhance membrane surface area, but also spatially sequester several integral membrane proteins. Proteomic analyses of microvillar membrane fractions have isolated numerous transmembrane transporters and channels involved in nutrient processing and transport^44^.

The recent revelation that epithelial microvilli are mechanosensitive presents an exciting new avenue to understanding luminal sensation during development, homeostasis and disease. However, no microvilli-specific mechanosensitive channels have yet been localized. Here, we used two distinct approaches to determine that TMC4 and 5 localize to the tips of microvilli in intestinal epithelial cells, presenting the first viable candidates for a mechanosensitive channel. First, we determined the tip localization of TMC4 in human IEC microvilli and the enrichment of TMC5 in mouse microvilli by Western blot. Second, we generated knockin mice expressing GFP-tagged TMC4 and mCherry-tagged TMC5 to confirm that both proteins localize to the distal tips of enterocyte microvilli.

Aside from their shared localization at the tips of actin protrusions, TMC4 and 5 share additional intriguing similarities with TMC1 and 2, the putative pore forming components of mechanosensitive channels at the tips of stereocilia in sensory cells of the inner ear. Adjacent stereocilia are connected by extracellular cadherin-based tip-links that are tensed when stereocilia are deflected and transmit forces to gate the TMC1 and 2-containing transduction channel. Microvilli are also interconnected via cadherin-based links^11^. Additionally, microvillar tips are the sites for secretion of numerous glycoproteins and glycolipids that form the glycocalyx (Fig. 5 and 6). We showed, using intravital microscopy, that the glycocalyx is subjected to luminal forces, which are likely transmitted to the proteins within the microvillar tip complex^45^. Another similarity involves lipid scrambling. TMC1 has been shown to be needed for lipid scrambling in stereocilia^46^, and here, we predict that TMC5 can be a lipid scramblase. Finally, the TMC1 and 2 stereocilia transduction channel complex requires CIB family proteins to function. We find that CIB3 is also present at intestinal microvillar tips, that it colocalizes with TMC4 and 5 in heterologous cells, and that it interacts directly with TMC5-IL1 in *in vitro* NMR spectroscopy experiments.

TMC4 and 5 are also predicted to have interesting differences with TMC1 and 2, which are believed to be cation-selective. Our homology modeling data predict that the pore in TMC4 and 5 is more positively charged than the TMC1 pore, and our AF2 models combined with simulations suggest that TMC4 and 5 may be more propense to permeate anions than TMC1 and 2 . This suggests that TMC4 and 5 act as anion or non-selective channels, which is consistent with a recent report describing a role for TMC4 as a Cl^-^ channel in papillae cells in the tongue^47^. Although this distinction suggests that TMC4 and 5 function is distinct from TMC1 and 2, it does not preclude their potential to be mechanosensitive channels. Anion efflux through ANO2 (also known as TMEM16B), a calcium activated chloride channel, has recently been established as an efficient mechanism of signal amplification in olfactory and gustatory receptor neurons^48^.

Overall, localization of TMC4 and 5 at the intestinal microvillar tips underscores similarities between microvilli and stereocilia. Based on these analogous TMC complexes, our modeling data, and the recent findings that microvilli act as mechanosensors^2,3^, we suggest that TMC4 and 5 are microvillar-mechanosensitive ion channels. Investigation of the contribution of mechanical stimuli to IEC microvilli regulation and function is relatively unexplored and these findings may have a significant impact on the understanding of luminal sensing in the gut.

## Methods

### Animals

All experimental procedures were conducted in accordance with the guide for the care and use of laboratory animals by NIH and approved by the animal care and use committees for the National Institute on Deafness and Other Communication Disorders (NIDCD ACUC, protocol #1215), and the University of Virginia Institutional Animal Care & Use Committee (UVA IACUC, protocol #4382). The following mice were used in this study:1) C57Bl/6, 2) TMC4-GFP, 3) TMC5-mCh and 4) TMC4-GFP: TMC5-mCh, 5) TMC1-mCherry^16^, 6) TMC2-GFP^16^. At least three animals were imaged per condition tested.

For fixed and frozen sample preparation, mice were euthanized by CO_2_ asphyxiation and then decapitated. Intestinal tissue was rapidly dissected and either directly frozen or fixed in 4% paraformaldehyde (PFA) in phosphate buffered saline (PBS) for downstream processing.

### Human tissue samples

Anonymized human intestinal tissue biopsies were acquired from the UVA Biorepository and Tissue Research Facility under the purview of the UVA Institutional Biosafety Committee (IBC number: 9987-22).

### Generation of knockin mice expressing TMC4-GFP and TMC5-mCherry

To label endogenous TMC4 and TMC5, CRISPR/Cas9 technology was implemented to knockin a GFP-tag at the 3’ end of *Tmc4* and a mCherry-tag at the 3’ end of *Tmc5*. A “linker-GFP” or “linker-mCherry cassette” was inserted into exon 15 of the Tmc4 gene or exon 22 of the Tmc5 gene as follows: 1) Cas9 mRNA and gRNA was produced by *in vitro* transcription; 2) the donor vector was constructed to contain a 5’ homologous arm (3.0 kb), the linker-fluorophore cassette, and a 3’ homologous arm (3.0 kb); 3) the mixture of Cas9 mRNA, gRNA and donor vector was microinjected into fertilized eggs from wild type C57BL/6J mice, and positive F0 mice were identified by PCR and sequencing; 4) F0 mice were crossed with wild type C57BL/6J mice to generate F1 mice. Mice were ultimately crossed to generate mice expressing both TMC4-GFP and TMC5-mCherry with the tags at the respective C-terminal ends.

### Antibodies

Affinity purified rabbit polyclonal antibody (pAb) against mouse TMC5 (epitope sequence: DFTVTHEKAVKLKQKNLSTE) was custom made by Covance. Anti-TMC4 (HPA048635) and anti-CIB3 (HPA043553) rabbit pAbs were purchased from Sigma. Anti-DDK mouse monoclonal antibodies (CAT#: TA150014) were purchased from Origene.

### Small Intestine Brush Border Isolation

The isolation procedure for brush borders was adapted from an established method^44^ and conducted as follows. The small intestine from 15 adult mice was dissected, immediately flushed with cold saline (150 mM NaCl, 2 mM imidazole), and immersed in a cold dissociation solution (200 mM sucrose, 12 mM EDTA, 18.9 mM KH2PO4, 78 mM Na2HPO4, pH 7.2) for 30 minutes. Isolated enterocytes in the solution were obtained using a Corning Stripette and subsequently washed three times by low-speed centrifugation (200 g for 10 min each) in fresh dissociation solution. The cell pellets were then resuspended in homogenization buffer (10 mM imidazole, 4 mM EDTA, 1 mM EGTA, 1 mM DTT, pH 7.2) supplemented with Complete Protease Inhibitor Cocktail (Sigma, 11836170001), and homogenized using a Fisher PowerGen 125 Homogenizer with four 15-second pulses. The homogenate was centrifuged at 1,000 g for 10 minutes to pellet the brush borders. The brush border pellets were washed several times in solution A (75 mM KCl, 10 mM Imidazole, 1 mM EGTA, 5 mM MgCl2, pH 7.2). To separate nuclei from intact brush borders, the pellets were resuspended in 50% sucrose solution A and carefully overlaid with 40% sucrose in Solution A. This gradient was centrifuged at 130,000 g for 1 hour at 4°C. The brush borders, located at the 40%/50% sucrose interface, were collected and washed three times with fresh solution A to remove residual sucrose.

### Western blot

Isolated brush border was homogenized at 4 °C for 30min in RIPA lysis buffer (G-Biosciences catalog #786-489) with cOmplete Protease Inhibitor Cocktail followed by centrifugation at 14,000g for 15 min. The resulting protein samples were prepared for gel electrophoresis by mixing them with an equal volume of 2x Laemmli Sample Buffer (Bio-rad, catalog #1610737) and boiling for 5 minutes. Protein concentrations were determined using a Bradford protein assay kit (Bio-rad, catalog #5000205), and 20 μg of protein was loaded into each lane of a 10-well SDS-PAGE gel. The protein sample was separated by SDS-PAGE and then transferred to a 0.22 μm nitrocellulose membrane (Bio-rad, catalog #1620097). The membrane was blocked in 5% skim milk overnight at 4 °C, and then the membrane was washed three times in 0.1% TBST followed by incubation with primary antibody diluted 1:1000 in 5% skim milk for 2 h at room temperature. After three washes in 0.1% TBST, the membrane was incubated with secondary antibodies (Jackson ImmunoResearch Secondary Antibodies) for 45 min. Following a final series of three washes in 0.1% TBST, the membrane was imaged using a LI-COR Odyssey imaging system to visualize the protein bands.

### COS7 cell transfection and immunofluorescence staining

COS7 cells (ATCC CRL-1651) were plated on coverslips and maintained at 37 °C in DMEM supplemented with 10% fetal bovine serum. Cells were transfected using Lipofectamine 3000 Transfection Reagent (Thermo Fisher) and incubated for 24 h. Samples were then fixed for 20 min in 4% formaldehyde in PBS, permeabilized for 30 min in 0.5% Triton X-100 in PBS and blocked at 4 °C in 10% normal goat serum (NGS) for 4 hours. Cells were then incubated in primary antibody (anti-TMC4, anti-TMC5 or anti-DDK, at 1 μg ml^−1^ in NGS) for 2 h, rinsed with PBS three times, stained with goat anti-rabbit, or goat anti-mouse secondary antibodies (Molecular Probes) conjugated with AlexaFluor-488 (A32731 or A32723) diluted 1:600 in PBS-NGS for 1 h, and counterstained with AlexaFluor-647 phalloidin (Molecular Probes, A-22287) diluted 1:600 in PBS-NGS. Cells were then rinsed with PBS three times and mounted using ProLong Gold Antifade reagent (Invitrogen). Confocal imaging of cell-culture experiments was performed using a TiE inverted fluorescence microscope (Nikon Instruments) equipped with a CSU-W1 Confocal Scanner Unit (Yokogawa), ORCA-Fusion BT Digital CMOS camera (Hamamatsu) and Apo TIRF 1.49 NA.

### Cryosection and immunofluorescence of intestinal tissue

After fixation in 4 % PFA in PBS for ∼ 30 min, intestinal tissues were put through a sucrose gradient from 10 %, to 20 % to 30 %, for cryoprotection. Once the tissue sank in the 30 % sucrose solution, it was placed in a mold, in optimal cutting temperature compound (O.C.T) and frozen on dry ice. Cryosections (10 µm thick) were then cut and adhered to silanated microscope slides. For whole-mount intestinal tissue preparation, intestine was extracted rinsed with PBS by perfusing through the lumen with syringe. The tissues were then fixed and cut into small segments that only contained the apical villus layer. For immunolabelling, sections and whole-mount intestinal tissues were first washed with 1X PBS pH 7.4, then permeabilized in 0.5 % Triton-X100 in PBS for 20 min and blocked in 10 % Normal Goat Serum (ThermoFisher Scientific, 50062Z) for ∼ 2 h at room temperature. Tissue was then incubated in primary antibody for 2 h before being rinsed with PBS three times. After this, tissue was incubated in goat anti-rabbit secondary antibodies conjugated with Alexa Fluor™ 568 (ThermoFisher Scientific, A-11011) diluted 1:1000, and Alexa Fluor™ Plus 405 Phalloidin (ThermoFisher Scientific, A-30104) diluted 1:200 in 10 % normal goat serum. Tissue was then rinsed with PBS three times and mounted using ProLong Gold Antifade reagent (Invitrogen) with a #1.5 coverslip.

### Spinning disc microscopes and imaging parameters

Microscopy was performed using Nikon TiE inverted fluorescence microscopes equipped with Yokogawa spinning disk confocal systems and either an Andor DU-888 camera or ORCA-Fusion BT Digital CMOS camera (Hamamatsu), and Apo TIRF 1.49 NA objectives. Sampling frequency at the camera sensor was 30 - 60 nm, enabled by a 1.5x tube lens and up to a total of 5x secondary magnification. NIS-Elements software (Nikon Instruments) was used to manage image acquisition.

### Transmission electron microscopy-freeze-substitution protocol

Freeze substitution was performed as previously described^49^. Fresh intestinal tissues were fine dissected in Gibco M199 and were directly frozen in the same media using a Life Cell CF-100 freezing machine. Frozen samples were freeze-substituted in a Leica Biosystems AFS with 1.5 % uranyl acetate in absolute methanol at −90 °C for 2 days, then infiltrated with HM20 Lowicryl resin (Electron Microscopy Sciences) over the course of 2 days at −45 °C. Finally, the resin was UV-polymerized for 3 days between −45 °C and 0 °C. Ultrathin sections (70 to 100 nm) were cut at and collected on hexagonal 300-mesh Ni grids (Electron Microscopy Sciences). Samples were imaged at zero electron energy loss on a 200-kV JEOL 2100 FX equipped with and an energy filter and a Gatan K2 Summit Direct Electron Detector camera. Acquisition software was DigitalMicrograph (Gatan); processing was done with DigitalMicrograph and FIJI and was limited to cropping and linear adjustments to brightness and contrast.

### Fast freezing and freeze-etch electron microscopy

Preparation of fixed samples: Samples were incubated with 2% glutaraldehyde overnight and then washed in ddH2O. Fixed samples were then fast frozen using a Life Cell CF-100 freezing machine. Frozen tissue was subjected to freeze-fracture at -110 °C in a Balzers freeze-fracture machine, followed by freeze-etch at -100 °C for 10 min. Freeze-etched samples were rotary shadowed with platinum and stabilized with carbon using electron-beam metal-evaporation guns (Cressington Scientific) to create replicas of the exposed surfaces. Replicas were cleaned with sodium hypochlorite, washed with ddH2O, and transferred onto 300 mesh hexagonal copper grids (Electron Microscopy Sciences).

### Data acquisition and electron tomography

Replicas were imaged on a JEOL 2100 electron microscope with an Orius 832 CCD camera (Gatan) or a OneView CMOS camera (Gatan). Single images were captured with DigitalMicrograph (Gatan). SerialEM was used to generate montages and acquire tilt series from -60° to + 60° at 1° increments. IMOD was used for montage blending and tomogram reconstruction^50^.

### Generation of homology-based structural models of TMC4 and 5

To identify the best structural template available, we used the Phyre2 server^51^. This server is updated weekly with the newest structures, and it would include the most recent structures of the TMEM16 and OSCA proteins. From all the templates identified, we selected those with the highest confidence percentage (>98%), coverage (>50% for TMC4 and >25% for TMC5), and sequence identity (8-17%). A total of 14 structural templates were identified for TMC5 and 11 for TMC4, all of them corresponding to cryo-EM and X-ray structures of several members of the TMEM16 and OSCA family of proteins. Therefore, identifying the best template was challenging since all the candidates presented a similar degree of coverage, sequence identity, and confidence. To include all the available template structures in our analysis, we generated structural models for each one of these templates and evaluated the models by several sequence-based (sequence coverage and identity) and structural-based parameters (local and global ProQM scores^52^ and Ramachandran plot statistics using PROCHECK^53^ to generate a model that best represent the structure of TMC4 and TMC5. Out of the three best models of TMC5 (PDB IDs: 4WIT, 5OC9, and 6E10), the model generated using the structure of the afTMEM16 (PDB ID: 6E10) as a template was selected as the best one. This model presented the highest ProQM score and the best Ramachandran plot statistics for a coverage with the second highest level of coverage of a 42.3% of the protein. In the case of TMC4, of the three best models (PDB IDs: 4WIT, 5OC9, and 6E10), the model based on the structure of the nhTMEM16 protein (PDB ID:4WIS) exceled over the other two, as it presented the highest ProQM score and most favorable Ramachandran plot statistics for the highest sequence coverage (84%). Sequence conservation of *Mus musculus* TMC4 or TMC5 proteins (UniProt ID: Q7Z404 and Q32NZ6, respectively) were analyzed using the ConSurf Server^54^. Homology sequences with a significance E-value of 10^-4^ and a sequence identity ranging from 35% to 95% were identified through HMMER^55^ after three iterations against the UniRef90 database. The closest 250 sequences were aligned using the MAFFT^56^ server to generate the multiple sequence alignment used to calculate conservation profiles of TMC4 and TMC5 proteins. The conservation score of each residue were mapped using a color-blind friendly code onto the TMC4 and TMC5. Protein structures were viewed with UCSF Chimera v1.12^57^. Final figures were generated in UCSF Chimera using the persistence of vision raytracer (POV-Ray) software v3.6.

### Generation of AF2-based structural models of TMC4 and 5

We used AF2^36^ for the predictions of mouse TMC4 and five dimeric complexes with and without CIB3. Sequences were obtained from NCBI for TMC4 (NP_861541.2), TMC5 (XP_011240226.1), and CIB3 (NP_001398710.1). Initially 50 predictions were generated for each monomer of TMC4 and TMC5. The predicted confidence values for the N-terminal ends of both proteins were low and thus predictions for the dimeric complexes were made with the N-terminal ends cut out from the models (residues 1-69 for TMC4 and 1-304 for TMC5). Similarly, we did 50 predictions for each dimeric complex using AF2 multimer and the top five best structures in each case were selected based on AF2 pTM scores. The predictions were visually inspected for anomalies using VMD^58^ and one of the top 5 predictions was selected for each of the complexes to be used in MD simulations.

### Preparation of simulation systems

The final models from the AF2 predictions (without N-termini as indicated above) were embedded in a POPC bilayer membrane using CHARMM-GUI^59^. The resultant system was solvated with explicit TIP3P water molecules and 300 mM NaCl while keeping systems neutral for electrostatic calculations using the particle mesh Ewald (PME) method. Systems including CIB3 and Ca^2+^ were predicted using AF2 and the shortened sequences of TMC4 and 5 along with the full-length sequence of CIB3. Then the CIB3 crystal structure (PDB:6WUD)^23^ was aligned to the CIB3 region of the AF2 predictions and the coordinates of the crystallographic Mg^2+^ ions were used to add Ca^2+^ to the AF2 predictions. These systems were then similarly embedded in a hydrated POPC bilayer and 300 mM NaCl. The overall system sizes for all the different complexes comprised ∼400,000 atoms (Table S1).

### NAMD simulations parameters and procedures

All-atom MD simulations^60^ were performed with CHARMM36 force field parameters for proteins and lipids and CMAP corrections^61–63^ using NAMD 2.13 in the Owens and Pitzer clusters at the Ohio Supercomputer Center. An integration time step of 2 fs was used with the SHAKE algorithm for constraining hydrogen atoms. Long-range electrostatic interactions were calculated using the PME method with grid density greater than 1 Å^-3^ and periodic boundary conditions. For all electrostatic and van der Waals calculations a 12 Å cutoff was used with a switching function starting at 10 Å. The Langevin thermostat was used to maintain a constant temperature of 310 K during production runs (damping coefficient of 0.1 ps^-1^ was used except for the equilibration part where the damping coefficient was 1 ps^-1^). A hybrid Nosé-Hoover Langevin piston method was employed to maintain a constant pressure of 1 atm (period 200 fs; damping timescale 50 fs). Atomic coordinates were saved every 5 ps.

To get a disordered bilayer membrane, the systems were first energy minimized for 2,000 steps and equilibrated for 0.5 ns with only the lipid tails free to move. This was followed by a further 2,000 steps minimization and 0.5 ns equilibration with constraints on all backbone atoms of the proteins. Final equilibration was done with no constraints for 5 ns, after which the production runs were started. For simulations with voltages, a constant electric field was applied to all atoms^41^ . TMD simulations^42^ were performed to slowly move channels from closed to open pore conformations. Pore backbone atoms were flagged and pulled using a spring constant of 200 Kcal/mol/Å^2^. Open pore conformations were locked by either fixing or constraining (*k* = 1 Kcal/mol/Å^2^) backbone atoms of the pore forming helices.

### Anton2 simulation parameters and procedures

Anton2 was used to perform 120-ns long equilibrations followed by 480-ns long simulations with voltage for systems TMC4, TMC4-CIB3, and TMC4-CIB3-Ca^2+^ (Simulations SA1l, SA1m, SA3l, SA3m, SA4l, and SA4m in Tables S1 and S2). Similarly, Anton 2 was used to perform 480-ns long simulations with voltage for systems TMC5, TMC5-CIB3, and TMC5-CIB3-Ca^2+^ (Simulations SA2k, SA5k, and SA6k in Tables S1 and S2)^64^. Anton2 compatible dms files were generated using the coordinate, velocity, and extended system files from NAMD equilibrated systems using the convertNAMDtoDMS.py script provided by the Pittsburgh Supercomputing Center. CHARMM36 forcefields were used and simulations were performed in the *NpT* ensemble (310 K, 1 atm) using the multigrator integration framework^65^ with a 2.5 fs integration timestep. Coordinates were saved every 120 ps. The MTK barostat and Nosé-Hoover thermostat were updated every 480 and 24 steps respectively. Semi-isotropic pressure coupling was used. A 12 Å cut-off was used for the calculations of Van der Waals interactions. A constant electric field was applied to all atoms to mimic transmembrane voltages^41^.

### MD simulation analyses

Average water density maps were calculated using the VolMap plugin in VMD^58^. Average electrostatic potential maps were calculated using PME and an Ewald factor of 0.25 and grid density > 1 Å^-3^. Both average water density and electrostatic maps were calculated using coordinates saved every 25 ps after the protein was aligned to the starting structure using backbone coordinates. Ionic currents were calculated by counting the number of ions that crossed the membrane bilayer from one side to the other through the pore. Number of ion crossings and the lipid crossing analysis were performed using in-house tcl scripts. Molecular images were rendered using VMD^58^, plots for electrostatic maps were generated using MATLAB, and ion conduction plots were generated using Xmgrace.

### Cloning and protein production, purification, and refolding for CIB3 and TMC5-IL1

DNA sequences encoding for *hs* CIB3 and *mm* TMC5-IL1 (equivalent to *hs* TMC5-IL1 T566A) were subcloned into *NdeI* and *XhoI* sites of the pET21a vector. All DNA constructs were sequence verified. Protein fragments were expressed in *Escherichia coli* BL21 Rosetta (DE3) (Novagen) or BL21 CodonPlus (DE3)-RIPL (Agilent) cells, which were cultured in LB or TB media, induced at OD_600_ ∼0.4-0.6 with 1 mM IPTG or 0.2 mM IPTG and grown at 30°C or 25°C for ∼16 h. Protein fragments for NMR were produced using the same cells cultured in M9 minimal media supplemented with MEM vitamin solution (Gibco) and 1 g/L ^15^NH_4_Cl (Cambridge Isotopes) to produce uniformly labeled [U-^15^N] protein. Cells were lysed by sonication in denaturing buffer (20 mM Tris HCl, pH 7.5, 6 M guanidine hydrochloride, 10 mM CaCl_2_, and 20 mM imidazole). The cleared lysates were loaded onto Ni-Sepharose (GE Healthcare) and eluted with denaturing buffer supplemented with 500 mM imidazole. Denatured proteins were (co-)refolded using MWCO 2,000 membranes (Spectra/Por 7) in dialysis buffer (400 mM Arg-HCl, 20 mM Tris-HCl, pH 7.5, 150 mM KCl, 50 mM NaCl, 2 mM CaCl_2_, 2 mM DTT) for ∼16 hr. Before starting refolding reactions, elution solutions were diluted to ∼0.5 mg/mL using the denaturing buffer, and then reduced by using 2 mM DTT in the diluted sample. Refolded and soluble proteins were concentrated using Vivaspin 20 centrifugal concentrators (10 kDa and 5 kDa molecular weight cutoff) and further purified on Superdex S75 columns (GE Healthcare) in SEC buffer (50 mM HEPES, pH 7.5, 126 mM KCl, 10 mM NaCl, 3 mM CaCl_2_, 5 mM DTT). Protein samples were concentrated down to a ∼0.5 mL volume for nuclear magnetic resonance (NMR) experiments.

### NMR experiments

Purified samples of *hs* [U-^15^N] CIB3 co-refolded in the presence of *hs* TMC5-IL1 T566A were concentrated to 180 μM in SEC buffer (50 mM HEPES pH 7.5, 3 mM CaCl_2_, 126 mM KCl, 10 mM NaOH, 5 mM DTT). The sample was then supplemented with 10% D_2_O (v/v) and 0.2 mM 2,2-dimethyl-2-silapentane-5-sulphonate (DSS) for field-frequency locking and chemical shift referencing. ^1^H-^15^N TROSY-HSQC spectra^66^ were recorded using a Bruker Avance III HD 800 MHz spectrometer equipped with a 5 mm TXI cryoprobe with z-gradients. The spectra were collected using 28 scans, 2048 x 256 complex points with a spectral width of 21.85 x 50.01 ppm in the ^1^H and ^15^N dimensions respectively. Spectra were processed and analyzed using NMRFx (http://nmrfx.org)^67^.

## Acknowledgements

This research was supported by the NIH, NIDCD Intramural Research Program (Z01 DC 000002 and DC000096) and the Center for Cell and Membrane Physiology, School of Medicine, at the University of Virginia through a start-up grant to S.E. Some of the electron micrographs were obtained at the NIDCD Advanced Imaging facility, ZIC DC000081. We acknowledge Travis Harrison-Rawn for assistance with sequence analyses. Simulations were carried out at the Ohio Supercomputer Center (OSC grants PAS1037 and PAA0217 to M.S.) and the Pittsburgh Supercomputing Center (PSC Anton 2 grant MCB150024P to M.S.). Nuclear magnetic resonance data were acquired at the Ohio State University Campus Chemical Instrument Center. J.S.M. was supported by an OSU/NIH molecular biophysics training grant (T32GM144293).

## Author contributions

B.K. and S.E. conceptualized the study, wrote the manuscript, and made the figures for experimental data, with contributions from all other co-authors. W.S.Z. and G.H. conducted COS7 cell transfections, Western blots and pulldowns, fixation and cryosectioning of human and mouse intestinal tissue, and immunofluorescence on COS7 cells and whole-mount and cryosections of the intestinal tissues. W.W.S. and E.S.K. prepared and imaged replicas of the murine intestinal tract. E.S.K. prepared and imaged immunogold labelled thin-sections brush border. R.C. carried out COS7 cell transfection and cryosectioning, and immunofluorescence on whole-mount and cryosections of mouse intestinal tissues. S.M., W-H.W. and Y.A. did molecular cloning, protein expression and purification, as well as refolding along with SEC experiments.

S.M. prepared AF2 models and did MD simulations and analyses with assistance from W-H.W. A.B. generated TMC4 and 5 homology-based models. J.S.M. did NMR experiments. J.S.M and M.P.F. analyzed NMR data.

M.S. supervised MD simulation and protein work and assisted with design of DNA constructs as well as with planning of protein production and refolding strategies. S.M and M.S prepared figures and text for computational and protein work parts with contributions from all other co-authors.

## Supplementary Materials

**Figure S1:**
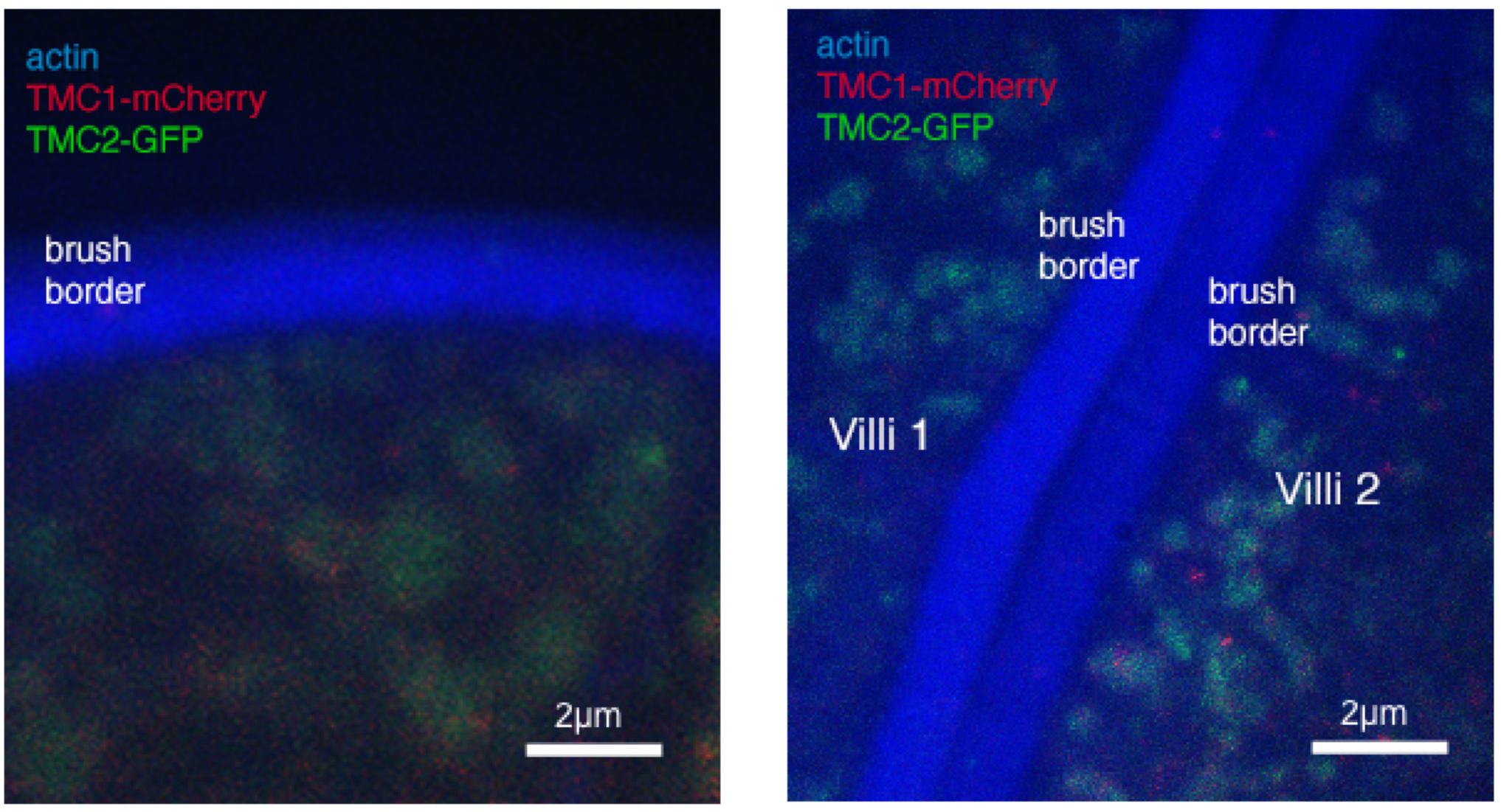
TMC1 and TMC2 are not expressed in IEC microvilli. Brush border microvilli stained with phalloidin (blue) in knockin mice expressing endogenous TMC2 tagged with GFP (green) and TMC1 tagged with mCherry (red)^1^.

**Figure S2:**
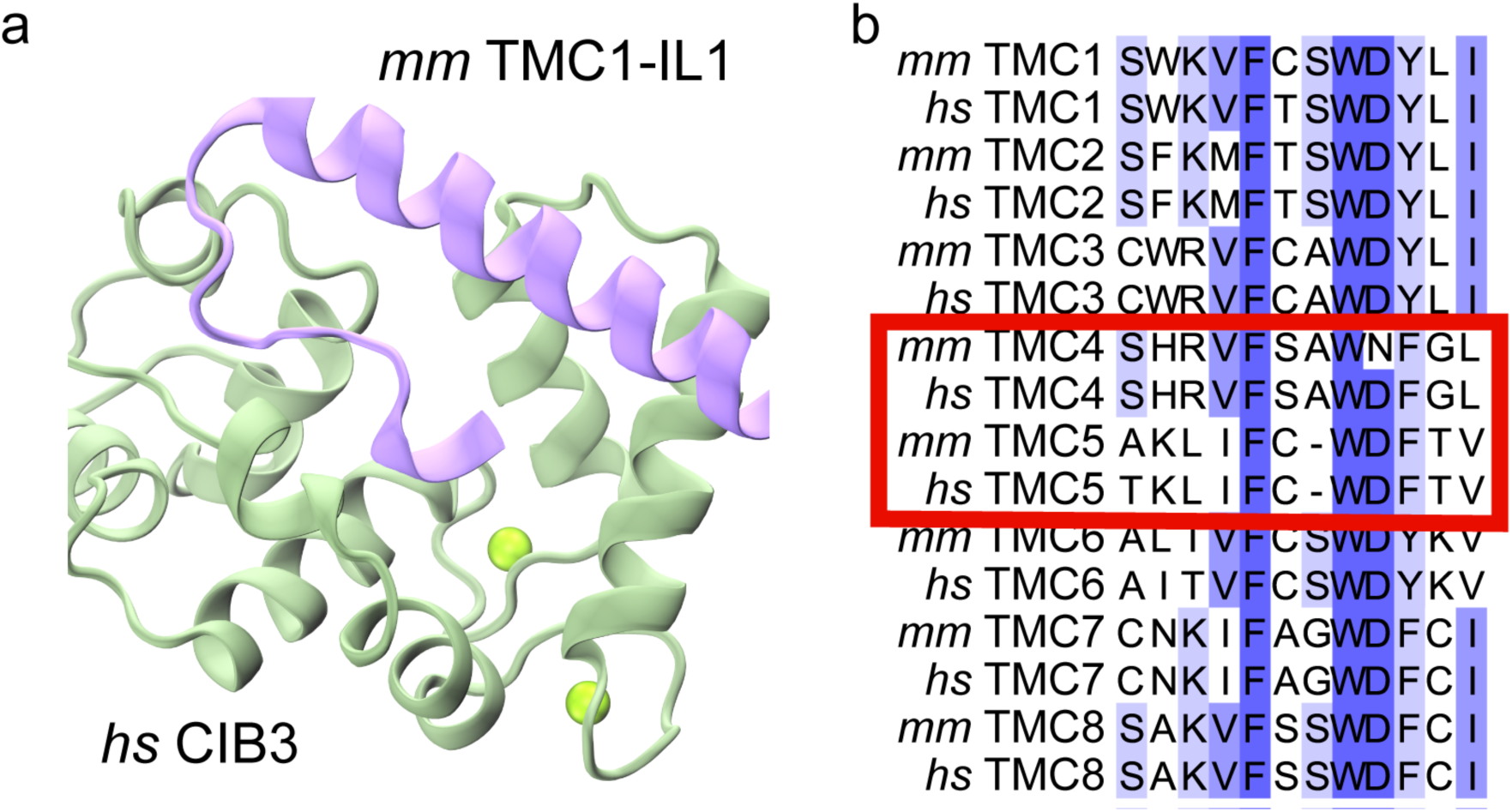
Conservation of TMC-IL1 and CIB interface. (a) Crystal structure of mouse (*Mus musculus*, *mm*) TMC1-IL1 and human (*Homo sapiens*, *hs*) CIB3 [PDB 6WUD]^2^. (b) Multiple sequence alignment comparing sequences of the IL1 region of mouse and human TMC1 through TMC8 (NP_083229.1, NP_619636.2, NP_619596.1, NP_542789.2, NP_808363.3, NP_001074001.1, NP_861541.2, NP_001138775.2, NP_001098722.1, NP_001098718.1, NP_663414.3, NP_001120670.1, NP_766064.2, NP_001287661.1, NP_001182017.1, NP_689681.2). The sequence motif FXXWDF/Y is conserved. Alignment is colored by sequence similarity with white being the lowest similarity and blue being the highest. Some columns might not be colored if deletions or changes to residue type are present (e.g., polar amino acid to hydrophobic).

**Figure S3.**
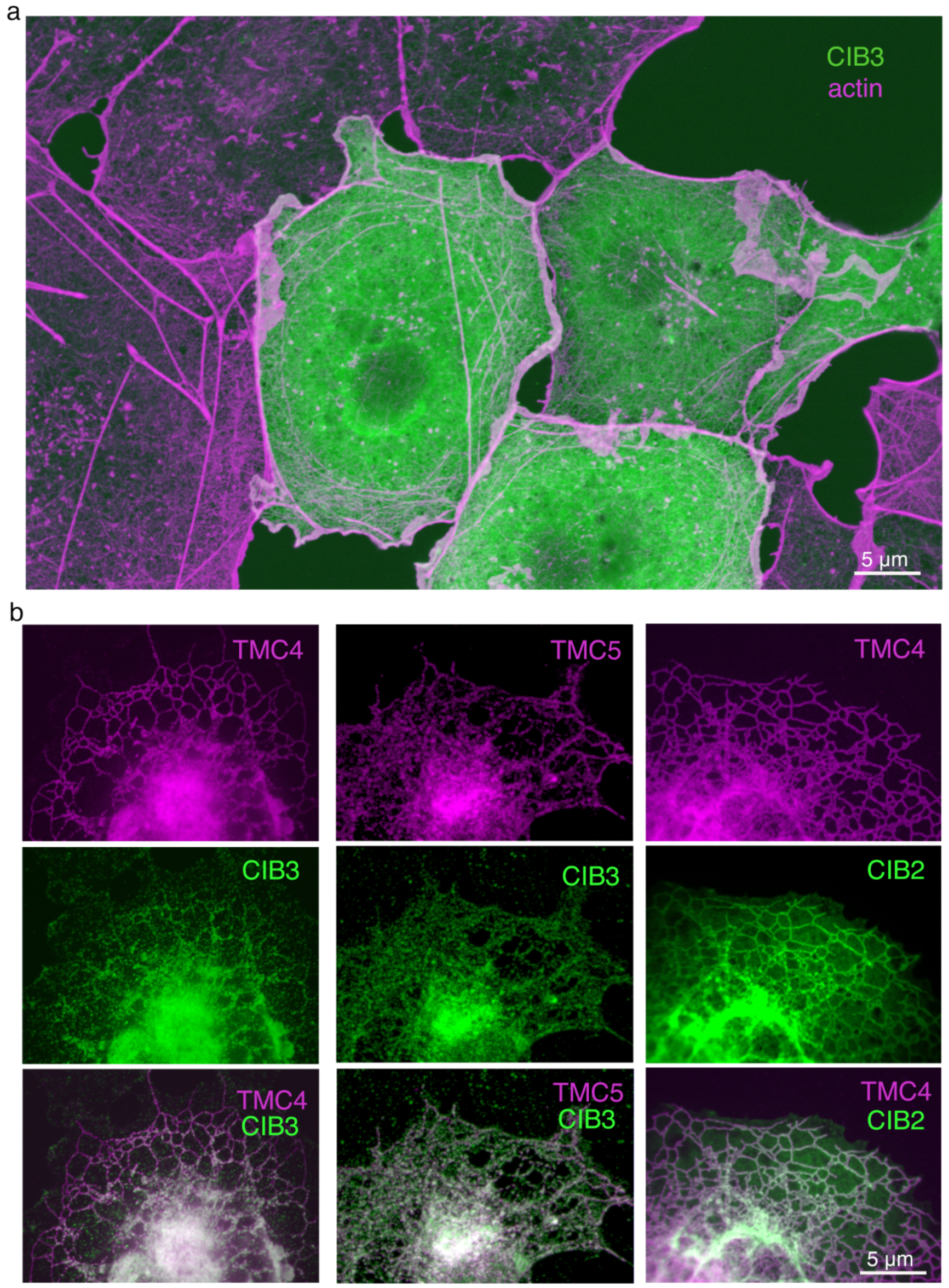
CIB family proteins colocalize with TMC4 and 5 in COS7 cells. (a) COS7 cells transfected with CIB3-DDK, labeled with anti-CIB3 antibody (green) and phalloidin to mark actin (magenta). (b) COS7 cells co-transfected with TMC4- or TMC5-mCherry (magenta) and CIB2- or CIB3-DDK, and immuno-labeled with anti-CIB3 antibody (green), or anti-DDK antibody for CIB2 (green).

**Figure S4.**
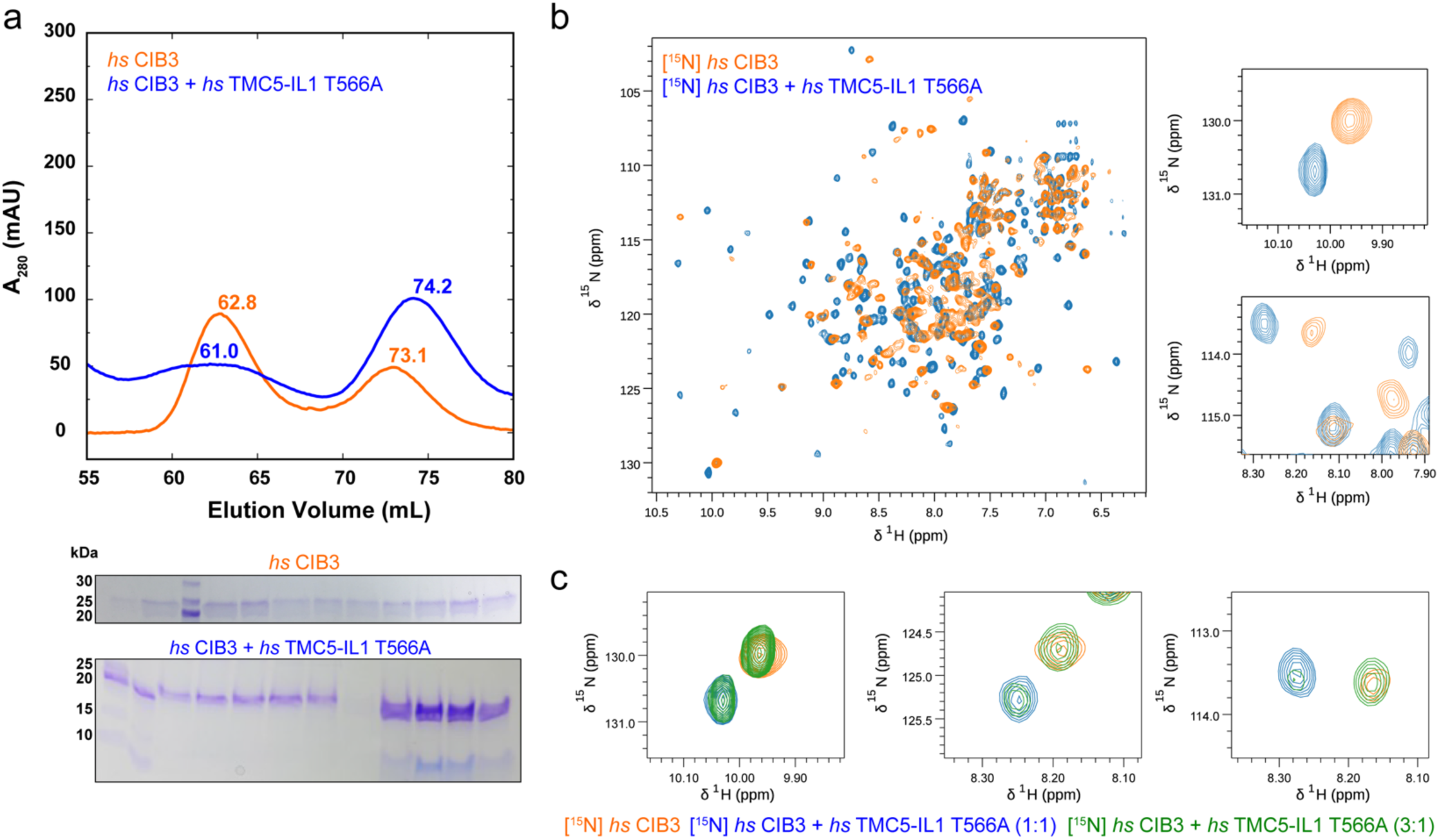
NMR data supports CIB3 and TMC5-IL1 complex formation. (a) Size exclusion chromatorgraphy (SEC) of CIB3 either refolded alone or co-refolded with *hs* TMC5-IL1 T566A (equivalent to *mm* TMC5-IL1). SEC curve and SDS-PAGE gel for CIB3 alone were adapted from Giese et al. (2024)^3^. (b) Overlay of ^1^H-^15^N TROSY-HSQC spectra of [U-^15^N]-CIB3 alone (orange) and bound to TMC5-IL1 (blue). (c) Detail of overlay of ^1^H-^15^N TROSY-HSQC spectra of [U-^15^N]-CIB3 alone (orange) and bound to TMC5-IL1 at 1:1 (blue) and 3:1 (green) CIB3:TMC5-IL1 ratios. Spectrum for [U-^15^N]-CIB3 alone was adapted from Giese et al. (2024)^3^.

**Figure S5.**
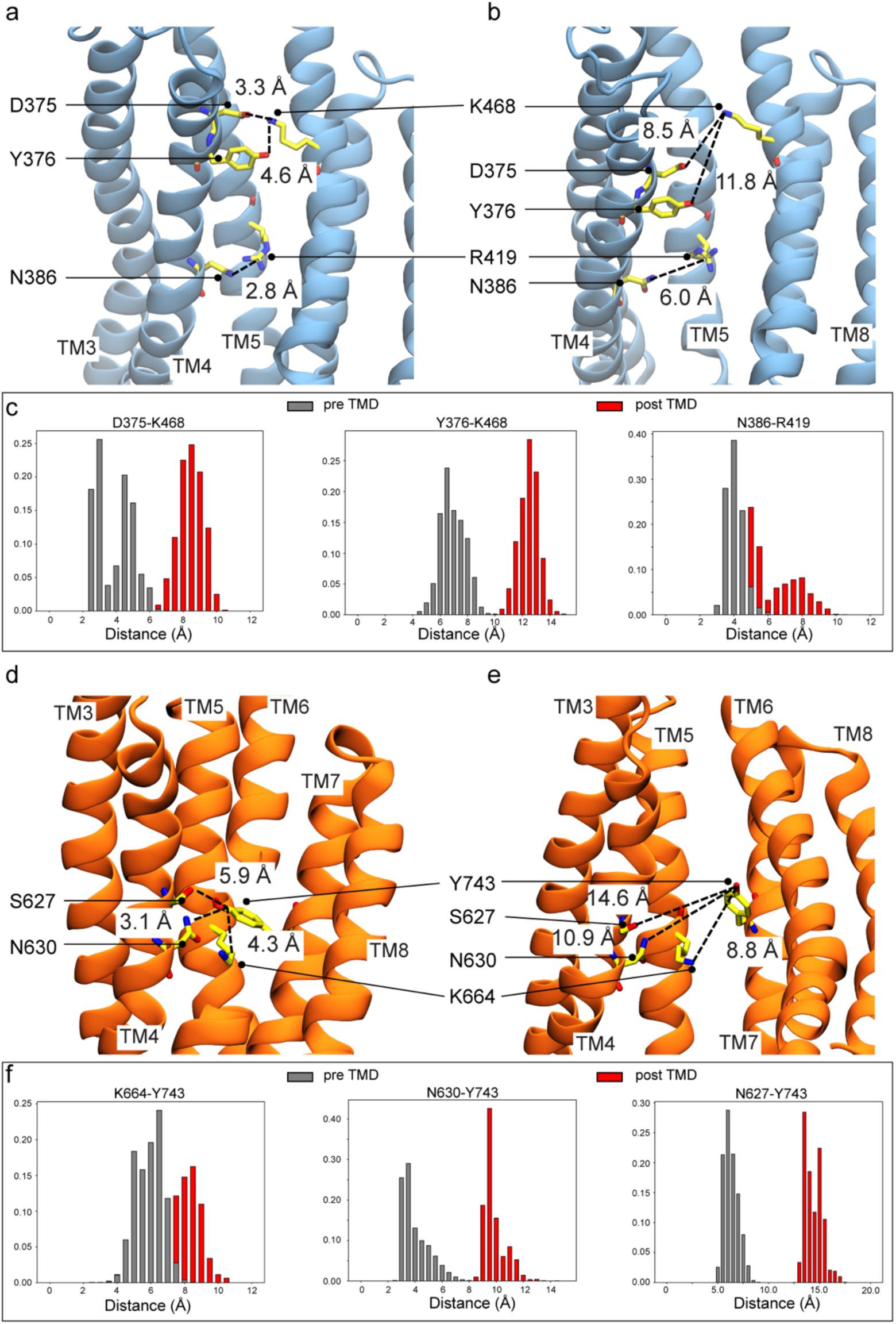
Closed and open pore models of TMC4 and 5. (a-b) AF2 and homology models representing closed and open pores for TMC4. Distances between D375 (TM4) – K468 (TM6), N386 (TM4) – R419 (TM5), and Y376 (TM4) – K468 (TM6) are shown to illustrate separation between TM4 and TM5/TM6 helices. (c) Distribution of distances between residues highlighted in (a) and (b) for the closed pore (pre TMD) and the open pore with constraints (post TMD). (d-e) AF2 and homology models representing closed and open pores for TMC5. Distances between S627 (TM4) – Y743 (TM7), N630 (TM4) – Y743 (TM7), and K664 (TM5) – Y743 (TM7) are shown to illustrate separation between TM7 and TM4/TM5 helices.

**Figure S6.**
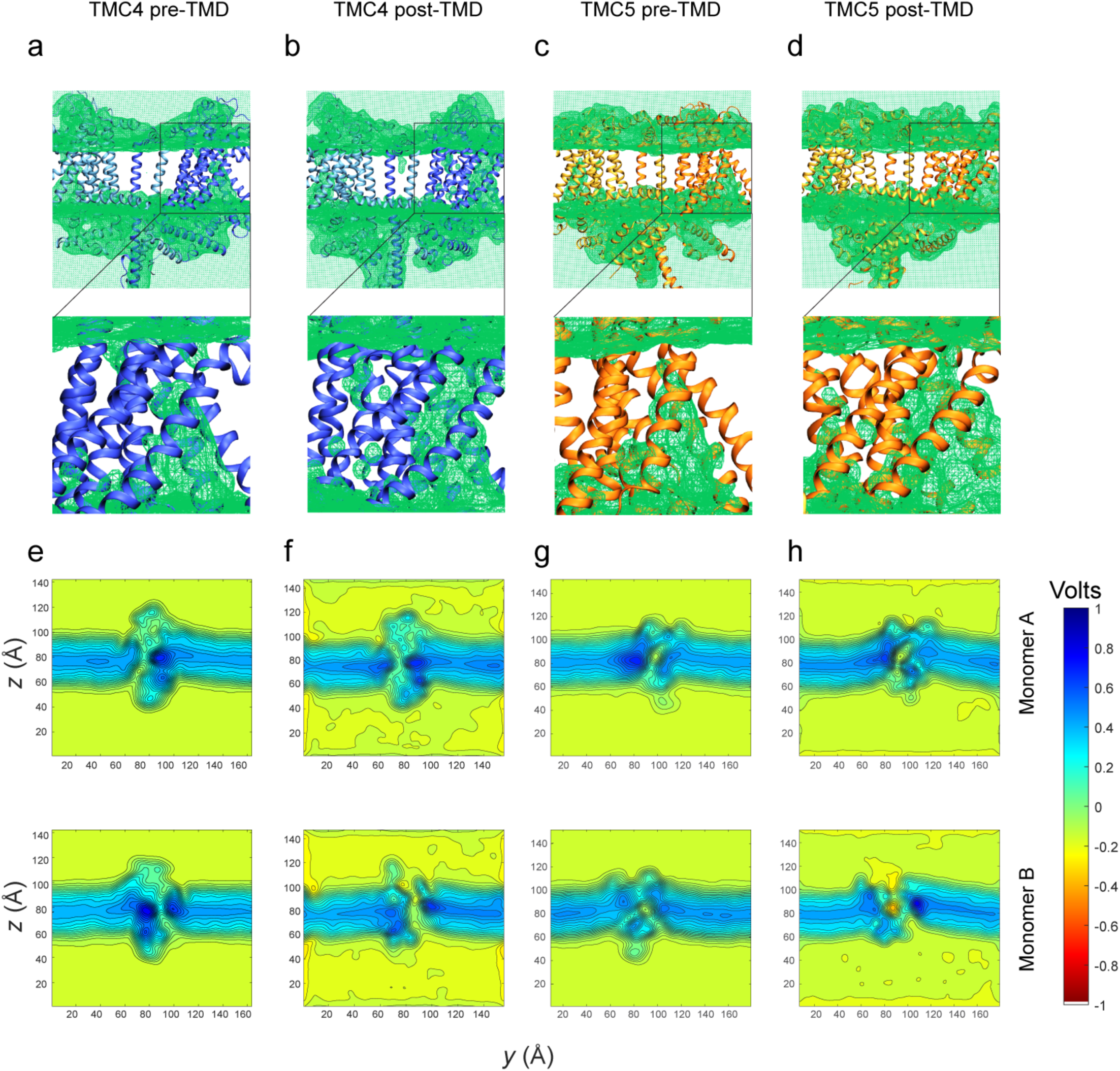
Water density and electrostatic maps for TMC4 and 5. (a-d) Water density maps for equilibration simulations of the TMC4 system prior (a) and post TMD (b), and of the TMC5 system prior (c) and post TMD (d). (e-f) Average electrostatic maps for TMC4 monomers A (top) and B (bottom), pre TMD (e) and post TMD equilibrium simulations. (g-h) Average electrostatic maps for TMC5 monomers A (top) and B (bottom) pre and post TMD (h). All post TMD simulations were performed with constraints on the backbone atoms of channel residues.

**Figure S7.**
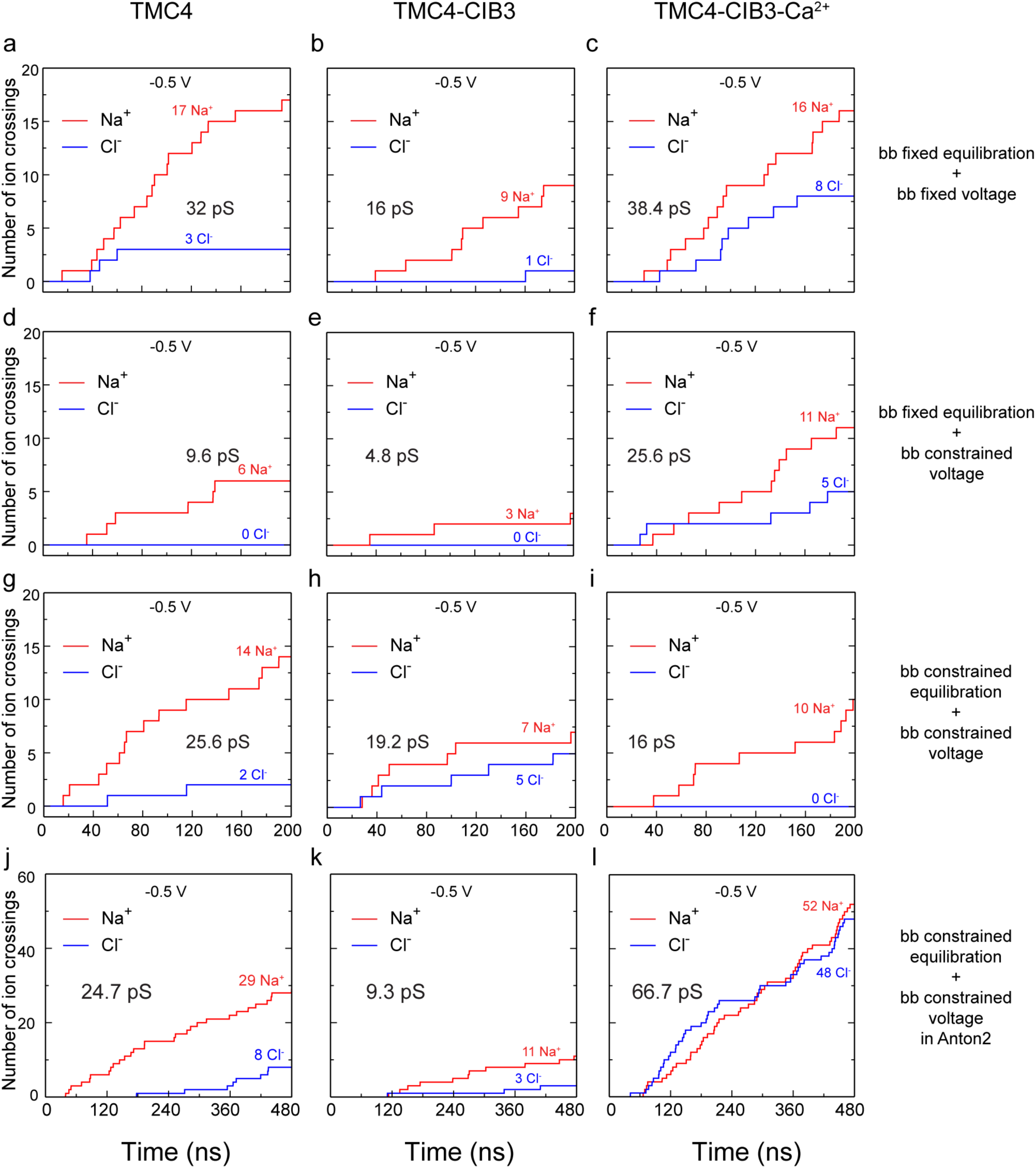
Number of ion crossings vs time for different simulations of TMC4, TMC4-CIB3, and TMC4-CIB3-Ca^2+^ systems. Number of ion crossings as a function of time for the conducting pore (chain B) of *mm* TMC4 after TMD was performed on chain B. Plots are shown for 200-ns long simulations at -0.5 V (a-i) and 480-ns long Anton2 simulations (j-l). Some panels are also in Fig. 9.

**Figure S8.**
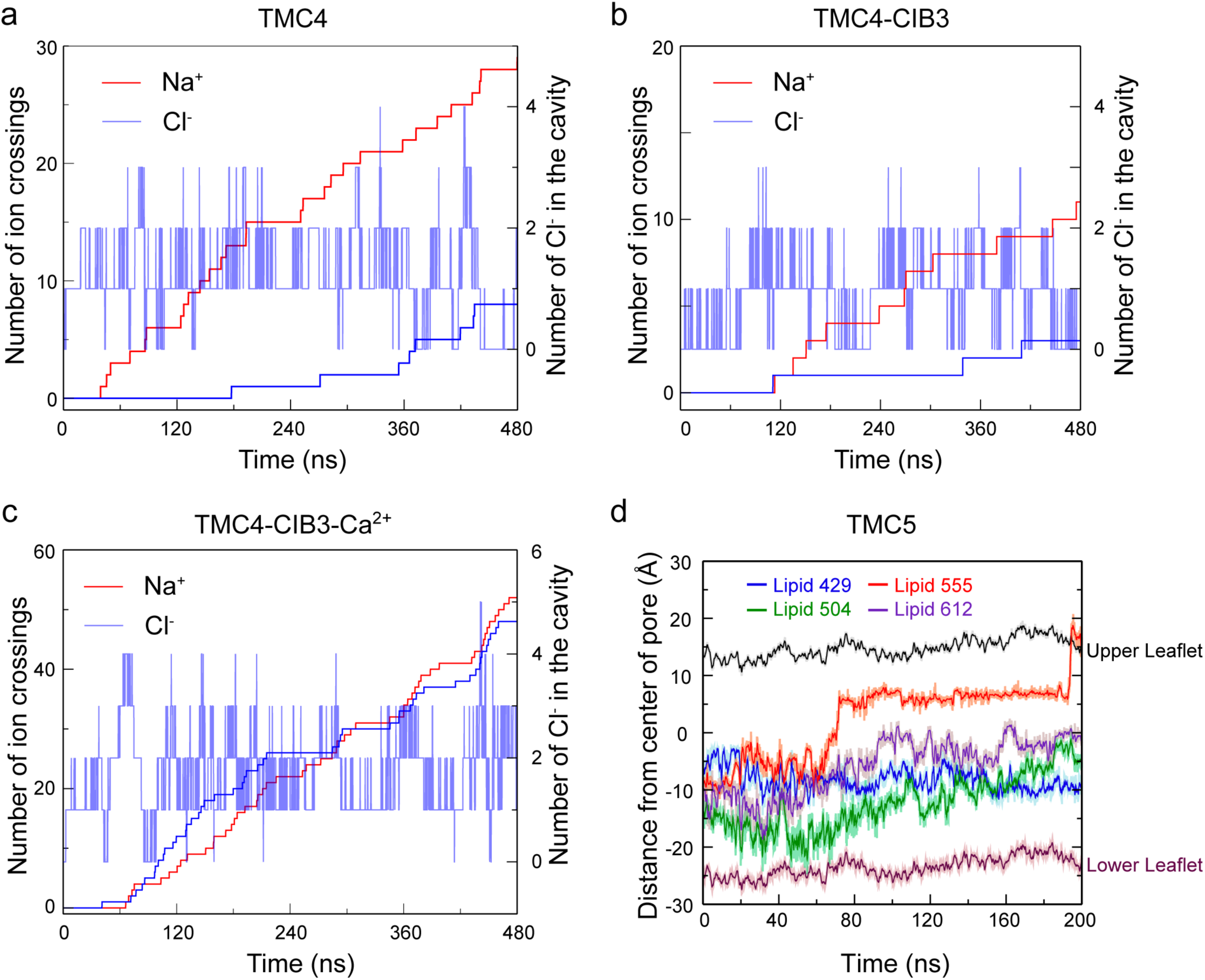
TMC4 and 5 pore properties. (a-c) Number of Na^+^ and Cl^-^ ions crossing and Cl^-^ ions present inside the cavity at a certain point of time from simulations SA1m, SA2m, and SA3m (Table S2). An ion was considered to be inside the cavity if it had a *z*-coordinate in between (C_cavity_ +7 Å) and (C_cavity_ -13 Å) where C_cavity_ = *z-*coordinate of center of the pore defined by residues 320 to 560. (d) Plot showing the position of P atoms of lipid headgroups as a function of time with respect to the center of the channel in TMC5 at -0.5 V. Lipid 555 (red) is moving from the intracellular side towards the extracellular side. This trajectory is also shown in Fig. 9m.

**Figure S9.**
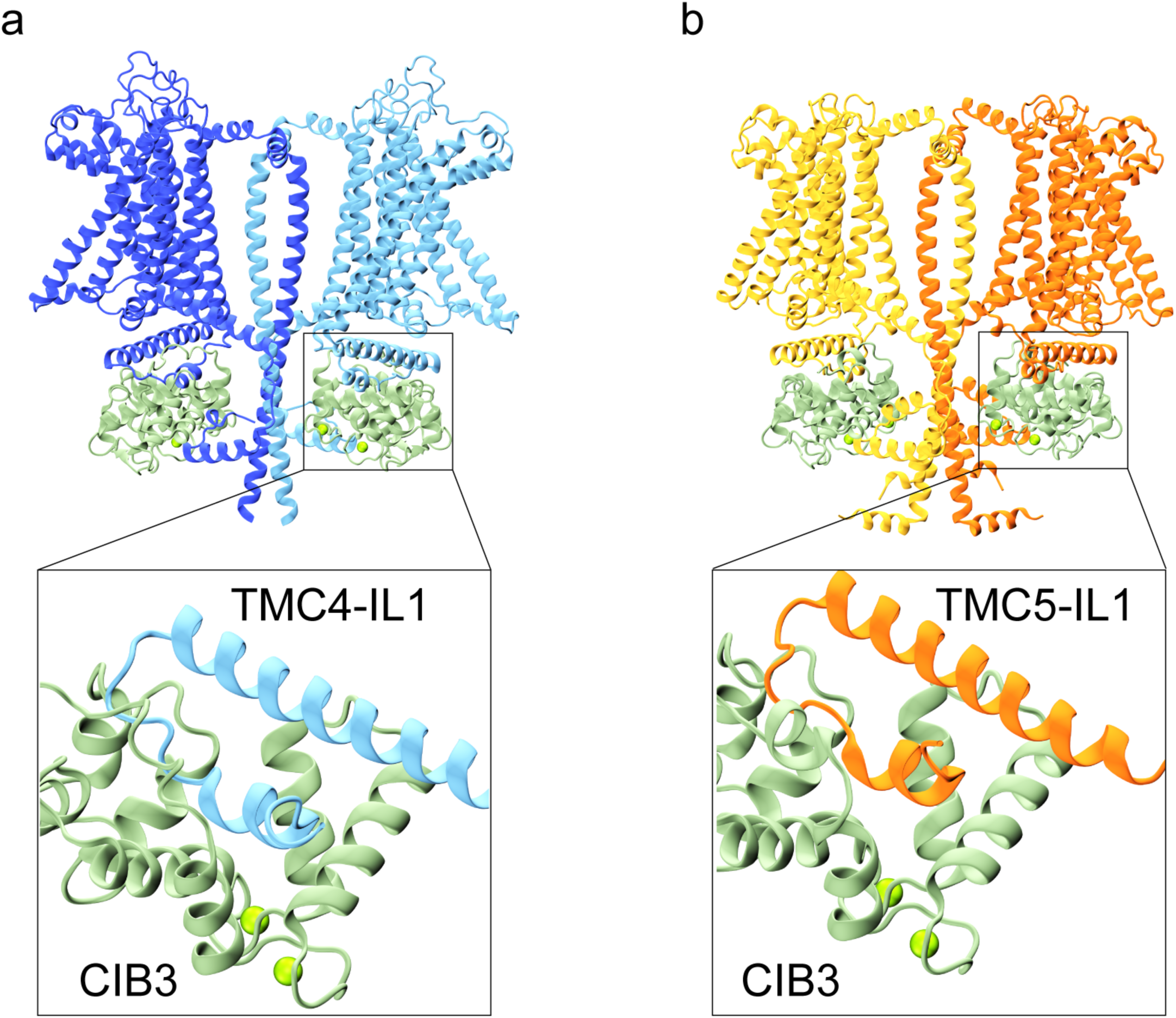
TMC and CIB interactions in AF2 models. (a-b) AF2 predicted models of CIB3 interacting with TMC4 (a) and 5 (b). Insets show details of the interaction with IL-1 for each protein. Ca^2+^ ions are shown as green spheres.

**Table S1.**
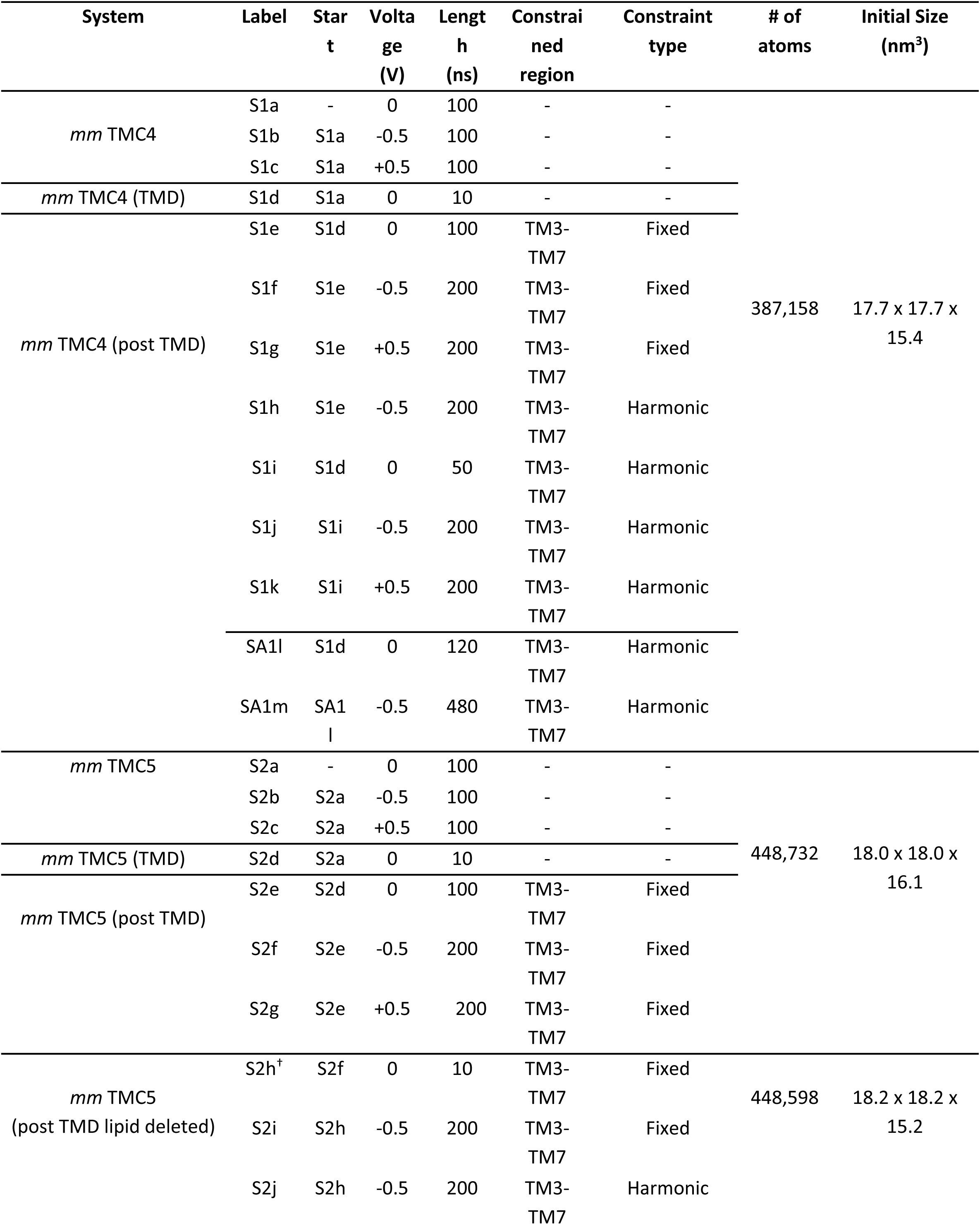
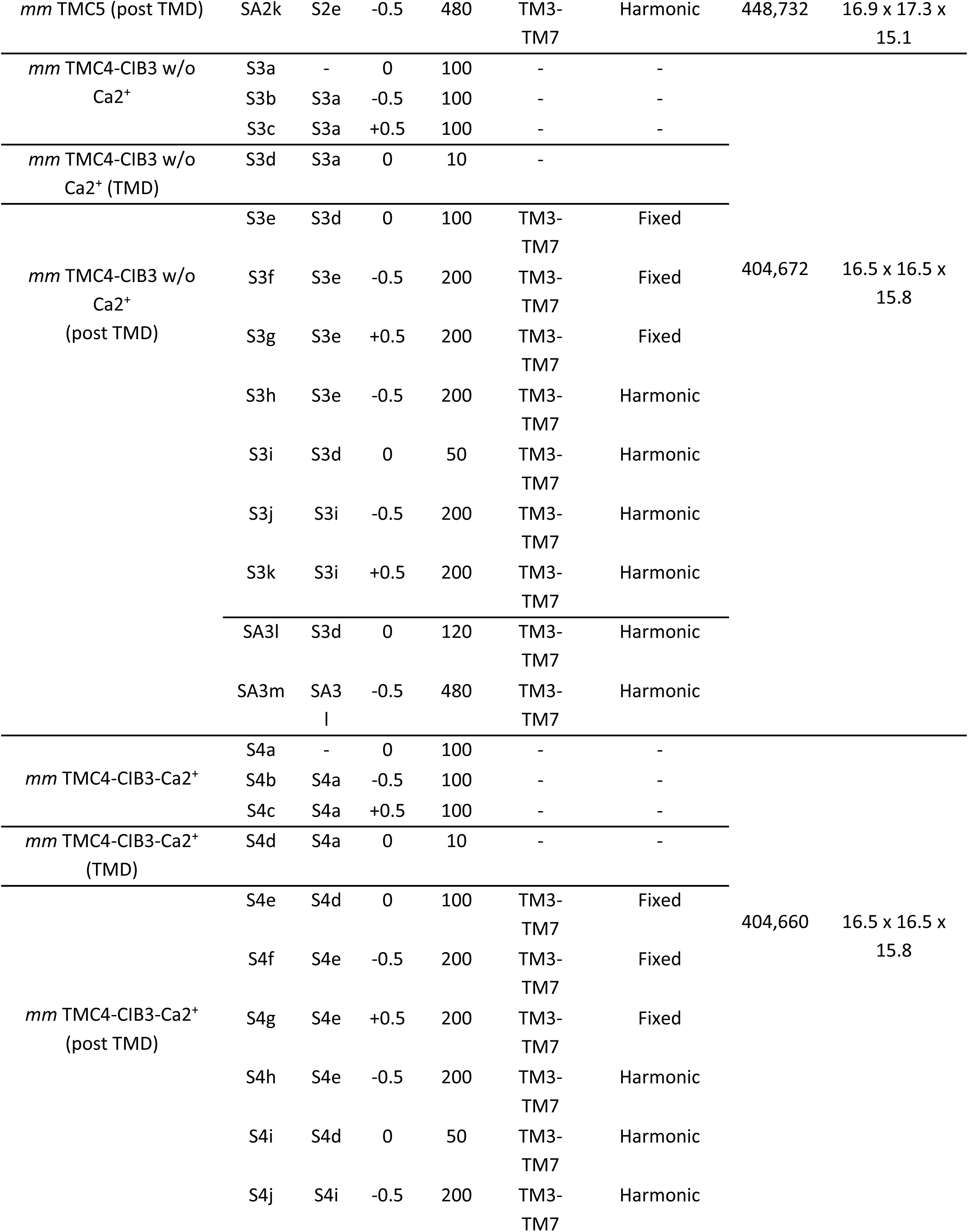
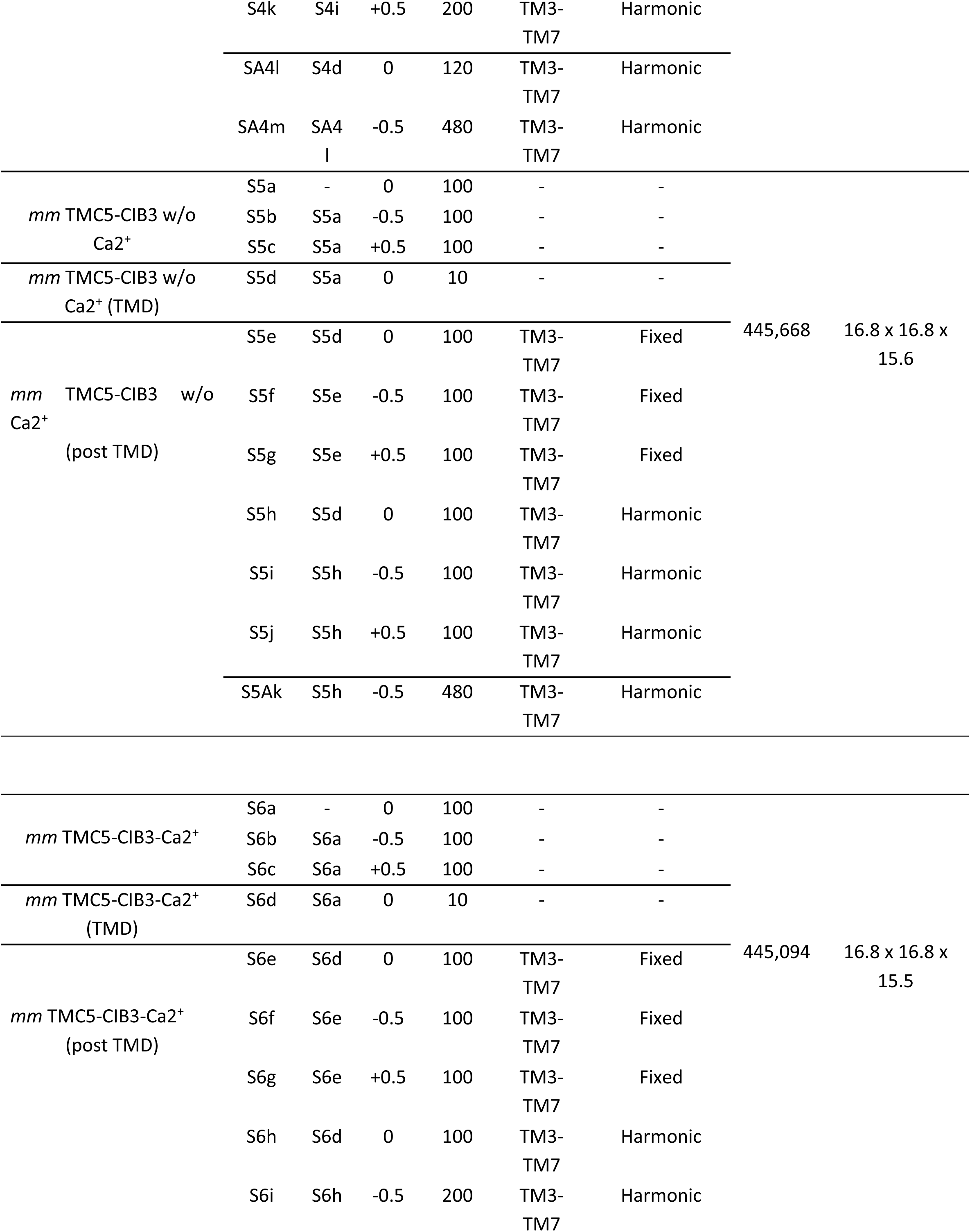
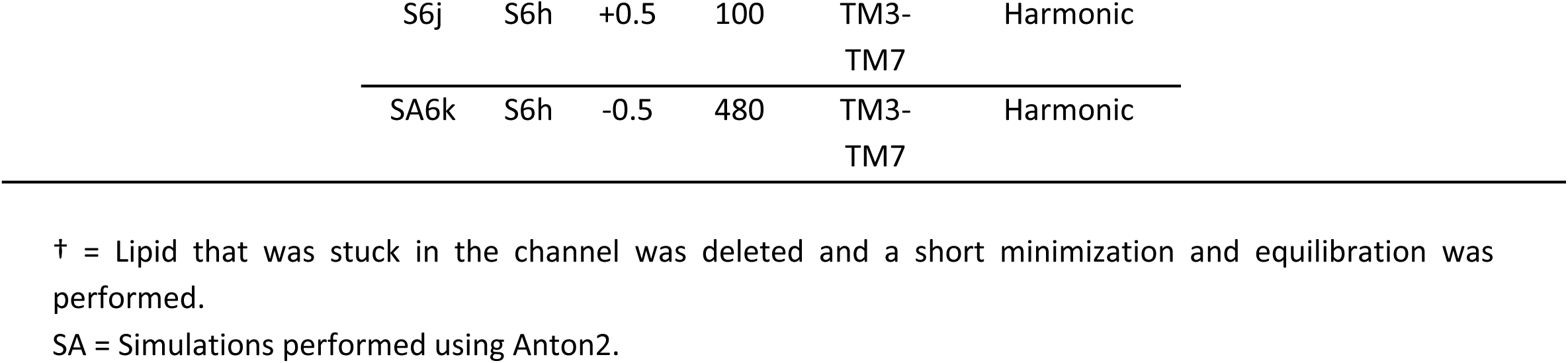
Summary of Simulations.

**Table S2.** Summary of Ion Conduction Events.

| Label | Voltage (V) | Length (ns) | Constrained region | Constraint type | Na+ (A) | Na+ (B) | Cl- (A) | Cl- (B) | I (pA) | C (pS) |
| --- | --- | --- | --- | --- | --- | --- | --- | --- | --- | --- |
| S1a | 0 | 100 | - | - | - | - | - | - | - | - |
| S1b | -0.5 | 100 | - | - | 0 | 0 | 0 | 0 | 0 | 0 |
| S1c | +0.5 | 100 | - | - | 0 | 0 | 0 | 0 | 0 | 0 |
| S1d | 0 | 10 | - | - | - | - | - | - | - | - |
| S1e | 0 | 100 | TM3-TM7 | Fixed | - | - | - | - | - | - |
| S1f | -0.5 | 200 | TM3-TM7 | Fixed | 0 | 17 | 0 | 3 | 16 | 32 |
| S1g | +0.5 | 200 | TM3-TM7 | Fixed | 0 | 2 | 0 | 3 | 4 | 8 |
| S1h | -0.5 | 200 | TM3-TM7 | Harmonic | 0 | 6 | 0 | 0 | 4.8 | 9.6 |
| S1i | 0 | 50 | TM3-TM7 | Harmonic | - | - | - | - | - | - |
| S1j | -0.5 | 200 | TM3-TM7 | Harmonic | 0 | 14 | 0 | 2 | 12.8 | 25.6 |
| S1k | +0.5 | 200 | TM3-TM7 | Harmonic | 0 | 4 | 0 | 0 | 3.2 | 6.4 |
| SA1l | 0 | 120 | TM3-TM7 | Harmonic | - | - | - | - | - | - |
| SA1m | -0.5 | 480 | TM3-TM7 | Harmonic | 0 | 29 | 0 | 8 | 12.3 | 24.7 |
| S2a | 0 | 100 | - | - | - | - | - | - | - | - |
| S2b | -0.5 | 100 | - | - | 0 | 0 | 0 | 0 | 0 | 0 |
| S2c | +0.5 | 100 | - | - | 0 | 0 | 0 | 0 | 0 | 0 |
| S2d | 0 | 10 | - | - | - | - | - | - | - | - |
| S2e | 0 | 100 | TM3-TM7 | Fixed | - | - | - | - | - | - |
| S2f | -0.5 | 200 | TM3-TM7 | Fixed | 0 | 0 | 0 | 0 | 0 | 0 |
| S2g | +0.5 | 200 | TM3-TM7 | Fixed | 0 | 0 | 0 | 0 | 0 | 0 |
| S2h<sup>†</sup> | 0 | 10 | TM3-TM7 | Fixed | - | - | - | - | - | - |
| S2i | -0.5 | 200 | TM3-TM7 | Fixed | 0 | 4 | 0 | 10 | 11.2 | 22.4 |
| S2j | -0.5 | 200 | TM3-TM7 | Harmonic | 0 | 7 | 0 | 12 | 15.2 | 30.4 |
| SA2k | -0.5 | 480 | TM3-TM7 | Harmonic | 0 | 0 | 0 | 0 | 0 | 0 |
| S3a | 0 | 100 | - | - | - | - | - | - | - | - |
| S3b | -0.5 | 100 | - | - | 0 | 0 | 0 | 0 | 0 | 0 |
| S3c | +0.5 | 100 | - | - | 0 | 0 | 0 | 0 | 0 | 0 |
| S3d | 0 | 10 | - | - | - | - | - | - | - | - |
| S3e | 0 | 100 | TM3-TM7 | Fixed | - | - | - | - | - | - |
| S3f | -0.5 | 200 | TM3-TM7 | Fixed | 0 | 9 | 0 | 1 | 8 | 16 |
| S3g | +0.5 | 200 | TM3-TM7 | Fixed | 0 | 1 | 0 | 0 | 0.8 | 1.6 |
| S3h | -0.5 | 200 | TM3-TM7 | Harmonic | 0 | 3 | 0 | 0 | 2.4 | 4.8 |
| S3i | 0 | 50 | TM3-TM7 | Harmonic | - | - | - | - | - | - |
| S3j | -0.5 | 200 | TM3-TM7 | Harmonic | 0 | 7 | 0 | 5 | 9.6 | 19.2 |
| S3k | +0.5 | 200 | TM3-TM7 | Harmonic | 0 | 2 | 0 | 3 | 4 | 8 |
| SA3l | 0 | 120 | TM3-TM7 | Harmonic | - | - | - | - | - | - |
| SA3m | -0.5 | 480 | TM3-TM7 | Harmonic | 0 | 11 | 0 | 3 | 4.7 | 9.3 |
| S4a | 0 | 100 | - | - | - | - | - | - | - | - |
| S4b | -0.5 | 100 | - | - | 0 | 0 | 0 | 0 | 0 | 0 |
| S4c | +0.5 | 100 | - | - | 0 | 0 | 0 | 0 | 0 | 0 |
| S4d | 0 | 10 | - | - | - | - | - | - | - | - |
| S4e | 0 | 100 | TM3-TM7 | Fixed | - | - | - | - | - | - |
| S4f | -0.5 | 200 | TM3-TM7 | Fixed | 0 | 16 | 0 | 8 | 19.2 | 38.4 |
| S4g | +0.5 | 200 | TM3-TM7 | Fixed | 0 | 0 | 0 | 0 | 0 | 0 |
| S4h | -0.5 | 200 | TM3-TM7 | Harmonic | 0 | 11 | 0 | 5 | 12.8 | 25.6 |
| S4i | 0 | 50 | TM3-TM7 | Harmonic | - | - | - | - | - | - |
| S4j | -0.5 | 200 | TM3-TM7 | Harmonic | 0 | 10 | 0 | 0 | 8 | 16 |
| S4k | +0.5 | 200 | TM3-TM7 | Harmonic | 0 | 0 | 0 | 0 | 0 | 0 |
| SA4l | 0 | 120 | TM3-TM7 | Harmonic | - | - | - | - | - | - |
| SA4m | -0.5 | 480 | TM3-TM7 | Harmonic | 0 | 52 | 0 | 48 | 33.3 | 66.7 |
| S5a | 0 | 100 | - | - | - | - | - | - | - | - |
| S5b | -0.5 | 100 | - | - | 0 | 0 | 0 | 0 | 0 | 0 |
| S5c | +0.5 | 100 | - | - | 0 | 0 | 0 | 0 | 0 | 0 |
| S5d | 0 | 10 | - | - | - | - | - | - | - | - |
| S5e | 0 | 100 | TM3-TM7 | Fixed | - | - | - | - | - | - |
| S5f | -0.5 | 100 | TM3-TM7 | Fixed | 0 | 0 | 0 | 0 | 0 | 0 |
| S5g | +0.5 | 100 | TM3-TM7 | Fixed | 0 | 0 | 0 | 0 | 0 | 0 |
| S5h | 0 | 100 | TM3-TM7 | Harmonic | - | - | - | - | - | - |
| S5i | -0.5 | 200 | TM3-TM7 | Harmonic | 0 | 0 | 0 | 0 | 0 | 0 |
| S5j | +0.5 | 100 | TM3-TM7 | Harmonic | 0 | 1 | 0 | 0 | 1.6 | 3.2 |
| SA5k | -0.5 | 480 | TM3-TM7 | Harmonic | 0 | 9 | 0 | 8 | 5.67 | 11.3<br>4 |
| S6a | 0 | 100 | - | - | - | - | - | - | - | - |
| S6b | -0.5 | 100 | - | - | 0 | 0 | 0 | 0 | 0 | 0 |
| S6c | +0.5 | 100 | - | - | 0 | 0 | 0 | 0 | 0 | 0 |
| S6d | 0 | 10 | - | - | - | - | - | - | - | - |
| S6e | 0 | 100 | TM3-TM7 | Fixed | - | - | - | - | - | - |
| S6f | -0.5 | 100 | TM3-TM7 | Fixed | 0 | 0 | 0 | 0 | 0 | 0 |
| S6g | +0.5 | 100 | TM3-TM7 | Fixed | 0 | 0 | 0 | 0 | 0 | 0 |
| S6h | 0 | 100 | TM3-TM7 | Harmonic | - | - | - | - | - | - |
| S6i | -0.5 | 200 | TM3-TM7 | Harmonic | 0 | 1 | 0 | 1 | 1.6 | 3.2 |
| S6j | +0.5 | 100 | TM3-TM7 | Harmonic | 0 | 0 | 0 | 0 | 0 | 0 |
| SA6k | -0.5 | 480 | TM3-TM7 | Harmonic | 0 | 0 | 0 | 0 | 0 | 0 |

## Movie Captions

**Movie 1.** mmTMC4-EQ.mp4. Chloride ions (green spheres) entering the TMC4 channel from the intracellular side during 100 ns equilibration simulation (Simulation S1a in Tables S1 and S2). Chain B of the protein is shown in light blue new cartoon representation in foreground while chain A has been shown in dark blue quicksurf representation. Chloride ions entering the channel oscillate among three basic residues R419 (TM5), R488 (TM6) and K549 (TM8) represented by yellow licorice with blue terminals (Nitrogen atoms). Water and lipid molecules have been omitted for visualization purposes.

**Movie 2.** mmTMC4-TMD.mp4. Targeted MD simulation showing the opening of chain B (light blue new cartoon) in TMC4 (Simulation S1d in Tables S1 and S2). TM4 slowly moves away from TM6, increasing the pore size. Polar residues are represented by yellow licorice. Blue quicksurf represents chain A. Water and lipid molecules have been omitted for visualization purposes.

**Movie 3.** mmTMC4-postTMD-conduction.mp4. Post-TMD ion crossings through chain B of TMC4, in the presence of -0.5 V (Simulation S1f in Tables S1 and S2). Chain B of TMC4 is shown using light blue new cartoon representation and chain A using blue quicksurf. Phosphorous atoms of lipid headgroups are shown using white spheres. Chloride ions (green spheres) cross from the intracellular to the extracellular side whereas sodium ions (yellow spheres) cross from the extracellular to the intracellular side. Lipid headgroups line up the periphery of the channel from both sides facilitating ion movement. Water molecules and lipid tails have been omitted for visualization purposes.

**Movie 4.** mmTMC5-lipid-flipping.mp4. Lipid flipping through chain B of TMC5 at -0.5 V (Simulations S2j in Tables S1 and S2). Chain B of TMC5 is shown using orange new cartoon representation in the foreground and chain A using dark yellow quicksurf representation. The phosphorous atoms of lipid headgroups are represented by white spheres. Lipid 555, represented by red spheres, undergoes flipping from intracellular to extracellular leaflet. Water, ions, and lipid tails have been omitted for visualization purposes.

**Movie 5.** mmTMC4-CIB3-Ca2+-conduction.mp4. Post-TMD ion crossings through chain B of the TMC4 in TMC4-CIB3-Ca2+ system at -0.5 V (Simulation S4f in Tables S1 and S2). Chain B of protein is shown using light blue new cartoon representation whereas chain A is shown using blue quicksurf. Phosphorous atoms of lipid headgroups are represented using white spheres. Chloride and sodium ions are represented using green and yellow spheres, respectively. We see more chloride ions crossing the channel in this system compared to only TMC4 (Movie 3). Water and lipid molecules have been omitted for visualization purposes.

## References

1. Delacour, D., Salomon, J., Robine, S. & Louvard, D. Plasticity of the brush border — the yin and yang of intestinal homeostasis. Nat. Rev. Gastroenterol. Hepatol. 13, 161–174 (2016).

2. Miura, S., Sato, K., Kato-Negishi, M., Teshima, T. & Takeuchi, S. Fluid shear triggers microvilli formation via mechanosensitive activation of TRPV6. Nat. Commun. 6, 8871 (2015).

3. Kim, S. W. et al. Shear stress induces noncanonical autophagy in intestinal epithelial monolayers. Mol. Biol. Cell 28, 3043–3056 (2017).

4. Müller, T., et al. *MYO5B* mutations cause microvillus inclusion disease and disrupt epithelial cell polarity. Nat. Genet. 40, 1163–1165 (2008).

5. Ruemmele, F. M. et al. Loss-of-function of MYO5B is the main cause of microvillus inclusion disease: 15 novel mutations and a CaCo-2 RNAi cell model. Hum. Mutat. 31, 544–551 (2010).

6. Burman, A. et al. A Novel mouse model derived from an uncharacterized MVID patient MYO5B mutation. Physiology 38, 5731773 (2023).

7. Shiner, M. & Birbeck, M. S. C. The microvilli of the small intestinal surface epithelium in coeliac disease and in idiopathic steatorrhoea. Gut 2, 277–284 (1961).

8. Bitner-Glindzicz, M. et al. A recessive contiguous gene deletion causing infantile hyperinsulinism, enteropathy and deafness identifies the Usher type 1C gene. Nat. Genet. 26, 56 (2000).

9. Hussain, K. et al. Infantile hyperinsulinism associated with enteropathy, deafness and renal tubulopathy: clinical manifestations of a syndrome caused by a contiguous gene deletion located on chromosome 11p. J. Pediatr. Endocrinol. Metab. JPEM 17, 1613–1621 (2004).

10. Gray, M. E. et al. Heterophilic and homophilic cadherin interactions in intestinal intermicrovillar links are species dependent. PLOS Biol. 19, e3001463 (2021).

11. Crawley, S. W. et al. Intestinal brush border assembly driven by protocadherin-based intermicrovillar adhesion. Cell 157, 433–446 (2014).

12. Grillet, N. et al. Harmonin Mutations Cause Mechanotransduction Defects in Cochlear Hair Cells. Neuron 62, 375–387 (2009).

13. Grati, M. & Kachar, B. Myosin VIIa and sans localization at stereocilia upper tip-link density implicates these Usher syndrome proteins in mechanotransduction. Proc. Natl. Acad. Sci. 108, 11476–11481 (2011).

14. Weck, M. L., Crawley, S. W., Stone, C. R. & Tyska, M. J. Myosin-7b Promotes Distal Tip Localization of the Intermicrovillar Adhesion Complex. Curr. Biol. 26, 2717–2728 (2016).

15. Crawley, S. W., Weck, M. L., Grega-Larson, N. E., Shifrin, D. A. & Tyska, M. J. ANKS4B Is Essential for Intermicrovillar Adhesion Complex Formation. Dev. Cell 36, 190–200 (2016).

16. Kurima, K. et al. TMC1 and TMC2 Localize at the Site of Mechanotransduction in Mammalian Inner Ear Hair Cell Stereocilia. Cell Rep. 12, 1606–1617 (2015).

17. Beurg, M. et al. Variable number of TMC1-dependent mechanotransducer channels underlie tonotopic conductance gradients in the cochlea. Nat. Commun. 9, 2185 (2018).

18. Pan, B. et al. TMC1 Forms the Pore of Mechanosensory Transduction Channels in Vertebrate Inner Ear Hair Cells. Neuron 99, 736–753.e6 (2018).

19. Ballesteros, A., Fenollar-Ferrer, C. & Swartz, K. J. Structural relationship between the putative hair cell mechanotransduction channel TMC1 and TMEM16 proteins. eLife 7,.

20. Medrano-Soto, A. et al. Bioinformatic characterization of the Anoctamin Superfamily of Ca2+-activated ion channels and lipid scramblases. PLoS ONE 13, e0192851 (2018).

21. Hahn, Y., Kim, D. S., Pastan, I. H. & Lee, B. Anoctamin and transmembrane channel-like proteins are evolutionarily related. Int. J. Mol. Med. 24, 51–55 (2009).

22. Giese, A. P. J. et al. Complexes of vertebrate TMC1/2 and CIB2/3 proteins form hair-cell mechanotransduction cation channels. eLife 12, (2023).

23. Liang, X. et al. CIB2 and CIB3 are auxiliary subunits of the mechanotransduction channel of hair cells. Neuron 109, 2131–2149.e15 (2021).

24. Giese, A. P. J. et al. CIB2 interacts with TMC1 and TMC2 and is essential for mechanotransduction in auditory hair cells. Nat. Commun. 8, 43 (2017).

25. Pontén, F., Jirström, K. & Uhlen, M. The Human Protein Atlas--a tool for pathology. J. Pathol. 216, 387–393 (2008).

26. Keresztes, G., Mutai, H. & Heller, S. TMC and EVER genes belong to a larger novel family, the TMC gene family encoding transmembrane proteins. BMC Genomics 4, 24 (2003).

27. Tarbell, J. M. & Pahakis, M. Y. Mechanotransduction and the glycocalyx. J. Intern. Med. 259, 339–350 (2006).

28. Jeong, H. et al. Structures of the TMC-1 complex illuminate mechanosensory transduction. Nature 610, 796–803 (2022).

29. Kamadurai, H. B. & Foster, M. P. DNA Recognition via Mutual-Induced Fit by the Core-Binding Domain of Bacteriophage λ Integrase. Biochemistry 46, 13939–13947 (2007).

30. Dyson, H. J. & Wright, P. E. Intrinsically unstructured proteins and their functions. Nat. Rev. Mol. Cell Biol. 6, 197–208 (2005).

31. Wright, P. E. & Dyson, H. J. Intrinsically unstructured proteins: re-assessing the protein structure-function paradigm. J. Mol. Biol. 293, 321–331 (1999).

32. Hahn, Y., Kim, D. S., Pastan, I. H. & Lee, B. Anoctamin and transmembrane channel-like proteins are evolutionarily related. Int. J. Mol. Med. 24, 51–55 (2009).

33. Clark, S., Jeong, H., Posert, R., Goehring, A. & Gouaux, E. The structure of the Caenorhabditis elegans TMC-2 complex suggests roles of lipid-mediated subunit contacts in mechanosensory transduction. Proc. Natl. Acad. Sci. U. S. A. 121, e2314096121 (2024).

34. Falzone, M. E. et al. Structural basis of Ca2+-dependent activation and lipid transport by a TMEM16 scramblase. eLife 8, e43229 (2019).

35. Brunner, J. D., Lim, N. K., Schenck, S., Duerst, A. & Dutzler, R. X-ray structure of a calcium-activated TMEM16 lipid scramblase. Nature 516, 207–212 (2014).

36. Jumper, J. et al. Highly accurate protein structure prediction with AlphaFold. Nature 596, 583–589 (2021).

37. Dang, S. et al. Cryo-EM structures of the TMEM16A calcium-activated chloride channel. Nature 552, 426–429 (2017).

38. Bushell, S. R. et al. The structural basis of lipid scrambling and inactivation in the endoplasmic reticulum scramblase TMEM16K. Nat. Commun. 10, 3956 (2019).

39. Alvadia, C. et al. Cryo-EM structures and functional characterization of the murine lipid scramblase TMEM16F. eLife 8, e44365 (2019).

40. Evans, R. et al. Protein complex prediction with AlphaFold-Multimer. 2021.10.04.463034 Preprint at 10.1101/2021.10.04.463034 (2022).

41. Gumbart, J., Khalili-Araghi, F., Sotomayor, M. & Roux, B. Constant electric field simulations of the membrane potential illustrated with simple systems. Biochim. Biophys. Acta 1818, 294–302 (2012).

42. Schlitter, J., Engels, M. & Krüger, P. Targeted molecular dynamics: a new approach for searching pathways of conformational transitions. J. Mol. Graph. 12, 84–89 (1994).

43. Fettiplace, R., Furness, D. N. & Beurg, M. The conductance and organization of the TMC1-containing mechanotransducer channel complex in auditory hair cells. Proc. Natl. Acad. Sci. 119, e2210849119 (2022).

44. McConnell, R. E., Benesh, A. E., Mao, S., Tabb, D. L. & Tyska, M. J. Proteomic analysis of the enterocyte brush border. Am. J. Physiol.-Gastrointest. Liver Physiol. 300, G914–G926 (2011).

45. Sun, W. W. et al. Nanoarchitecture and dynamics of the mouse enteric glycocalyx examined by freeze-etching electron tomography and intravital microscopy. *Commun*. Biol. 3, 5 (2020).

46. Ballesteros, A. & Swartz, K. J. Regulation of membrane homeostasis by TMC1 mechanoelectrical transduction channels is essential for hearing. 2021.09.24.461722 Preprint at 10.1101/2021.09.24.461722 (2021).

47. Kasahara, Y. et al. TMC4 is a novel chloride channel involved in high-concentration salt taste sensation. J. Physiol. Sci. 71, 23 (2021).

48. Dibattista, M., Pifferi, S., Hernandez-Clavijo, A. & Menini, A. The physiological roles of anoctamin2/TMEM16B and anoctamin1/TMEM16A in chemical senses. Cell Calcium 120, 102889 (2024).

49. Ebrahim, S. et al. Dynamic polyhedral actomyosin lattices remodel micron-scale curved membranes during exocytosis in live mice. Nat. Cell Biol. 21, 933–939 (2019).

50. Kremer, J. R., Mastronarde, D. N. & McIntosh, J. R. Computer visualization of three-dimensional image data using IMOD. J. Struct. Biol. 116, 71–76 (1996).

51. Kelley, L. A., Mezulis, S., Yates, C. M., Wass, M. N. & Sternberg, M. J. E. The Phyre2 web portal for protein modeling, prediction and analysis. Nat. Protoc. 10, 845–858 (2015).

52. Ray, A., Lindahl, E. & Wallner, B. Model quality assessment for membrane proteins. Bioinforma. Oxf. Engl. 26, 3067–3074 (2010).

53. Laskowski, R. A., MacArthur, M. W., Moss, D. S. & Thornton, J. M. PROCHECK: a program to check the stereochemical quality of protein structures. J. Appl. Crystallogr. 26, 283–291 (1993).

54. Ashkenazy, H. et al. ConSurf 2016: an improved methodology to estimate and visualize evolutionary conservation in macromolecules. Nucleic Acids Res. 44, W344–350 (2016).

55. Finn, R. D., Clements, J. & Eddy, S. R. HMMER web server: interactive sequence similarity searching. Nucleic Acids Res. 39, W29–37 (2011).

56. Katoh, K., Misawa, K., Kuma, K. & Miyata, T. MAFFT: a novel method for rapid multiple sequence alignment based on fast Fourier transform. Nucleic Acids Res. 30, 3059–3066 (2002).

57. Pettersen, E. F. et al. UCSF Chimera--a visualization system for exploratory research and analysis. J. Comput. Chem. 25, 1605–1612 (2004).

58. Humphrey, W., Dalke, A. & Schulten, K. VMD: visual molecular dynamics. J. Mol. Graph. 14, 33–38, 27–28 (1996).

59. Jo, S., Kim, T., Iyer, V. G. & Im, W. CHARMM-GUI: a web-based graphical user interface for CHARMM. J. Comput. Chem. 29, 1859–1865 (2008).

60. Karplus, M. & Petsko, G. A. Molecular dynamics simulations in biology. Nature 347, 631–639 (1990).

61. Klauda, J. B. et al. Update of the CHARMM all-atom additive force field for lipids: validation on six lipid types. J. Phys. Chem. B 114, 7830–7843 (2010).

62. Huang, J. & MacKerell, A. D. CHARMM36 all-atom additive protein force field: validation based on comparison to NMR data. J. Comput. Chem. 34, 2135–2145 (2013).

63. Buck, M., Bouguet-Bonnet, S., Pastor, R. W. & MacKerell, A. D. Importance of the CMAP correction to the CHARMM22 protein force field: dynamics of hen lysozyme. Biophys. J. 90, L36–38 (2006).

64. Shaw, D. E. et al. Anton 2: Raising the Bar for Performance and Programmability in a Special-Purpose Molecular Dynamics Supercomputer. in *SC ’14: Proceedings of the International Conference for High Performance Computing, Networking*, Storage and Analysis 41–53 (2014). doi:10.1109/SC.2014.9.

65. Lippert, R. A. et al. Accurate and efficient integration for molecular dynamics simulations at constant temperature and pressure. J. Chem. Phys. 139, 164106 (2013).

66. Nietlispach, D. Suppression of anti-TROSY lines in a sensitivity enhanced gradient selection TROSY scheme. J. Biomol. NMR 31, 161–166 (2005).

67. Norris, M., Fetler, B., Marchant, J. & Johnson, B. A. NMRFx Processor: a cross-platform NMR data processing program. J. Biomol. NMR 65, 205–216 (2016).

## References

1. Kurima, K. et al. TMC1 and TMC2 Localize at the Site of Mechanotransduction in Mammalian Inner Ear Hair Cell Stereocilia. Cell Rep. 12, 1606–1617 (2015).

2. Liang, X. et al. CIB2 and CIB3 are auxiliary subunits of the mechanotransduction channel of hair cells. Neuron 109, 2131–2149.e15 (2021).

3. Giese, A. P. J. et al. Complexes of vertebrate TMC1/2 and CIB2/3 proteins form hair-cell mechanotransduction cation channels. eLife 12, (2023).

